# Clonally Convergent Tph/Tfh-like CD4^+^ T Cells are Shared Across Recovery and Recurrence in Experimental Periodontitis

**DOI:** 10.64898/2026.11.04.760965

**Authors:** A Sharma, V Olmedo, L Rojas, S Suliman, S Mohammadi, K Lauwens, G Nguyen, K Cruz-Dwyer, A Aguilar Angulo, Y Khan, J Dang, C Bluff, N Yakar, S Tornaas, DE Costea, H Dongre, Z Guney, F Oner, B Bor, A Shamshad, H Hasturk, G Hajishengallis, L. K Kirkwood, A Kantarci, C Alvarez Rivas

## Abstract

Periodontitis is a chronic inflammatory disease of the tooth-supporting structures (periodontium) that requires life-long maintenance to prevent progression. Using a sensitized model of recurrent periodontitis in mice (SeRPD), in which regulatory breakdown was induced by temporary Treg depletion concurrent with disease, we mapped the gingival T cell imprint across each disease phase. Transient regulatory breakdown left a durable mucosal imprint after clinical recovery, characterized by enrichment of local and regional CD4⁺ T cells. Paired single-cell RNA and TCR sequencing of gingival immune cells identified a convergent clonotype expanded from recovery into recurrence that carried a Tfh/Tph-like transcriptional identity. An analogous CD4⁺ T cell population was detected in human periodontitis lesions by single-cell transcriptomics. Together, these findings support a paradigm in which clinical recovery does not ensure a local T cell repertoire reset; instead, a reservoir of antigen-experienced, tissue-adapted CD4⁺ T cells persist in the mucosa and participate in recurrence.

**Highlights:**

- The murine SeRPD model induced local and regional enrichment of antigen-experienced CD4^+^ T cells.
- A clonally expanded convergent clonotype was shared in recovery and recurrence phases.
- After clinical recovery, convergent CD4^+^ T clones expressed a Tfh/Tph-like program.
- Human periodontitis lesions were associated with an analogous Tfh/Tph-like population.

**In Brief:** Sharma et al. show that periodontitis accompanied by regulatory breakdown, leaves a lasting, clonally expanded Tfh/Tph-like population within recovered gingiva that is also present during recurrence. A comparable CD4^+^ T cell-population was associated with human periodontitis, suggesting a memory reservoir for relapse.

## Introduction

Mucosal tissue barriers are a point of interaction between microbial communities and the immune system (1). Due to this tight interface, the immunity within mucosae is tailored to the specific tissue needs. A critical component of mucosal immunity is its regulatory mechanisms, such as regulatory T cells (Tregs), which suppress uncontrolled inflammatory responses (2). However, under physiological and pathological conditions, temporary immune regulatory disruptions can occur due to external and internal factors such as acute stress, acute inflammatory reactions, trauma and hormonal fluctuations (3–6). Whether such regulatory disruptions leave a lasting imprint on the mucosal T cell compartment remains poorly understood.

The gingiva is the portion of the oral mucosa associated with the teeth, and part of the tooth-supporting structures known as periodontium (7). Periodontitis is a microbiome-driven, chronic inflammation of the periodontium that affects nearly 50% of the global adult population and is mechanistically linked to multiple inflammatory morbidities (8, 9). The condition is characterized by persistence, as diagnosed patients require lifelong maintenance to prevent progression (10, 11). The recurrent nature of the disease suggests the participation of organized recall responses not yet understood. It also provides a clinically relevant framework to investigate how coexisting disease and momentary regulatory breakdown affect T cell persistence upon recovery.

T cells are critical components of the adaptive immune response in the oral mucosa (12, 13). After antigen presentation, through the interaction of the T cell receptor (TCR) with peptide-MHC complexes, CD8^+^ and CD4^+^ T cells clonally expand and acquire effector functions (14). Among CD4^+^ T helper subtypes, T helper 17 (Th17) cells contribute to periodontitis pathogenesis through IL-17 production (15), while Foxp3-expressing Treg cells attenuate inflammation through multiple immune suppressive mechanisms, making their transient disturbance a tractable experimental tool to study long-term T cell priming (16). During chronic mucosal inflammation, specialized CD4^+^ T cells, such as T peripheral helper (Tph) cells, sustain localized immune hubs and recruit antibody-producing B cells by secreting IL-21. These cells remain locally retained and have been recognized as drivers of refractory, chronic mucosal inflammation (17). Unlike follicular helper T (Tfh) cells, which provide B cell help within secondary lymphoid organs, Tph cells operate at sites of extrafollicular inflammation, sharing B-cell help functions and express of inhibitory receptors PD-1 and ICOS, but lack CXCR5 expression and germinal center activity (18). Whether these specialized T cell types accumulate during recurrent mucosal inflammation following regulatory disruptions remains unclear.

Clonally expanded effector T cells have a relatively short lifespan and undergo a contraction phase mediated by apoptosis. A fraction of surviving cells differentiates into memory T cells, expressing a transcriptomic program that supports long-term survival and enables a rapid and enables a rapid, robust secondary response upon antigen re-encounter (19, 20). Additionally, some surviving T cells may also display an exhausted phenotype, characterized by upregulation of inhibitory receptors such as PD-1 and CTLA-4 (21). Local clonal selection determines the persistence of activated T cells and their tissue-adaptability (22), since not all microbially derived peptides are recognized by T cells in periodontitis. Earlier studies suggest that T cell clonotypes selectively expand in response to a limited number of antigens in periodontal lesions, a phenomenon known as oligoclonality (23).

To study the impact of periodontitis and transient regulatory breakdown in mucosal T cell compartments, we developed an experimental model that reproduced the recurrent nature of the human disease, characterized by sustained inflammation, prolonged antigenic stimulation, high T cell infiltration, tissue retention, and exhaustion after recovery (24). We used a sensitized mouse model of recurrent ligature-induced periodontitis in which temporary depletion of Tregs was followed by resolution of inflammation after ligature removal. Transient regulatory disruption during periodontitis led to a marked gingival retention of CD4⁺ antigen experienced T cells and the persistent, highly expanded clonotypes after recovery. Upon disease recurrence (secondary ligature application without regulatory disruption), we detected phenotypic and clonal convergence of gingival Tfh/Tph-like T cells, as revealed by integrated paired single-cell RNA sequencing (scRNA-seq) and T cell receptor sequencing (TCR-seq) analyses. The clonally expanded population shared key transcriptomic features with a periodontitis-associated CD4⁺ T cell cluster in human periodontitis. Taken together, these findings highlight the lasting tissue imprint of sensitized periodontitis, even after structural and regulatory resolution, and clonally persistent Tfh/Tph-like CD4⁺ T cells arise as a candidate for antigen-driven recall responses in recurrent mucosal inflammation.

## Results

### Human periodontitis is characterized by gingival enrichment of antigen-experienced and tissue-adapted CD4^+^ T cells

First, we confirmed the presence of antigen-experienced T cells in human gingival mucosa using imaging mass cytometry (IMC), the CD45RO marker for antigen experience was used in healthy gingival mucosa (GM) and gingiva from periodontal lesions (PD) **(Supplementary Table 1)**. Qualitative analysis of the IMC images showed that in GM, CD45RO⁺CD4⁺ T cells localized primarily to perivascular regions (defined by αSMA labeling), whereas in PD these cells were predominantly peri-epithelial (defined by ECAD labeling; **Figure 1A**). In PD, the density of CD45RO⁺CD4⁺ T cells per mm² was significantly higher than both CD45RO⁺CD8⁺ T cell density in PD (p = 0.021) and CD45RO⁺CD8⁺ T cell density in GM (p = 0.018) (**Figure 1B**). There were no significant differences between CD45RO⁺CD4⁺ T cells per mm² in PD compared to GM, and CD45RO⁺CD8⁺ T cell per mm² in PD compared to GM. In both conditions, most CD4⁺ T cells expressed CD45RO (61.3% in GM and 66.6% in PD; **Figure 1B**). Thus, in periodontitis, CD4^+^ T cells dominate the CD45RO^+^ compartment relative to CD8^+^ T cells in gingiva.

**Figure 1.**
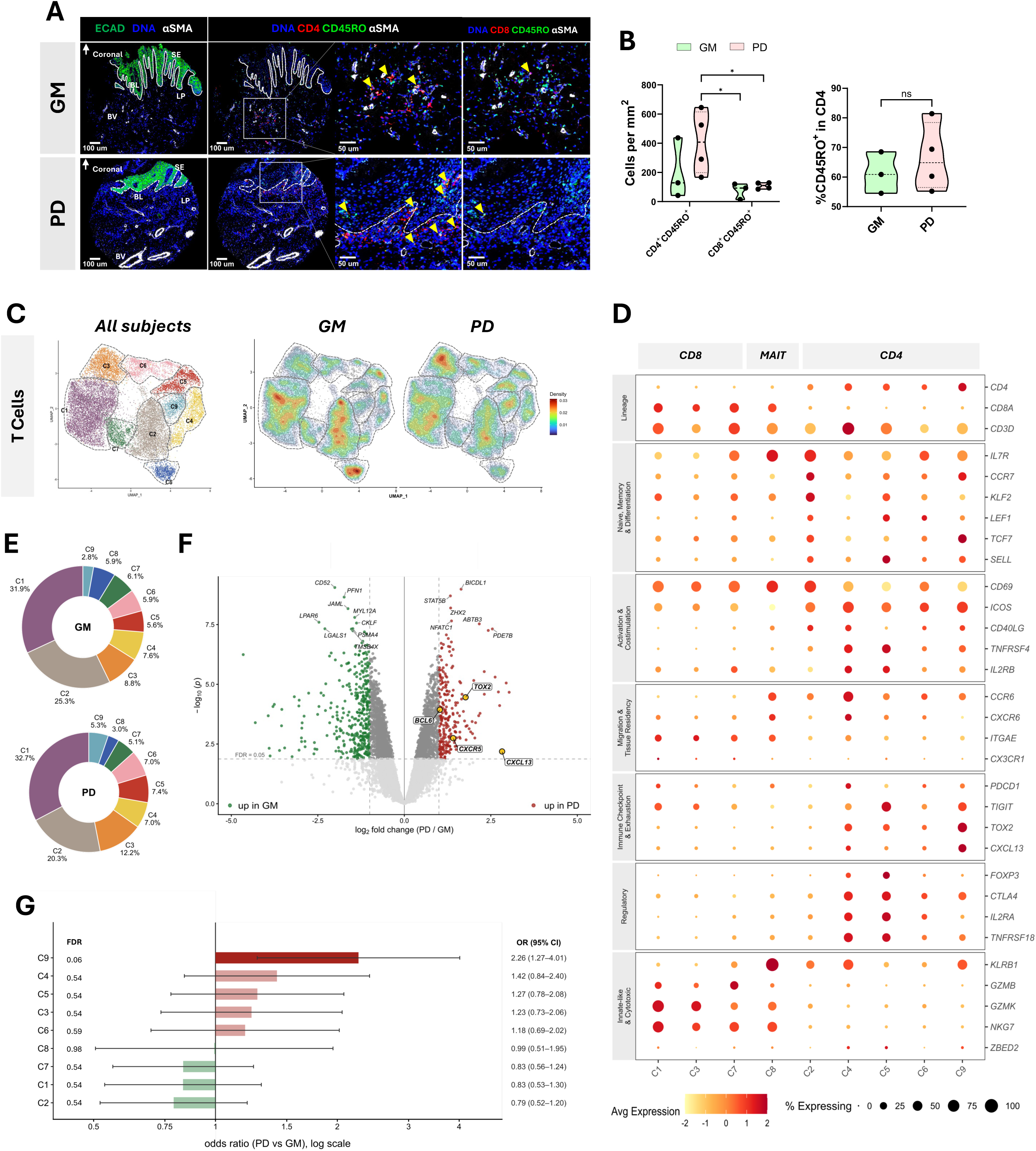
Human periodontitis is characterized by gingival enrichment of antigen-experienced and tissue-adapted CD4⁺ T cells. A) Imaging mass cytometry (IMC) images of normal gingival mucosa (GM) and periodontitis-affected gingival mucosa (PD). White arrowheads indicate coronal directionality relative to the teeth. Dashed white lines delineate the basal lamina (BL, ECAD⁺) and blood vessels (BV, αSMA⁺). Sulcular epithelium (SE) and lamina propria (LP) are annotated. The white square marks the region shown in the zoomed inset. Yellow arrows indicate CD4⁺CD45RO⁺ or CD8⁺CD45RO⁺ cells, identified by co-detection of red and green channels (yellow in the overlay). B) Quantification of IMC data across GM (n= 3 donors, one donor provided two samples) and PD (n= 4 donors) biopsies: density of CD4⁺CD45RO⁺ and CD8⁺CD45RO⁺ cells per mm² of tissue, and percentage of CD45RO⁺ cells among all CD4⁺ cells. Multiple comparisons for cell density were performed using two-way ANOVA with Fisher’s LSD test; the percentage comparison used the Mann-Whitney U test. Significance was set at p < 0.05 (*); ns = not significant. To characterize the T cell landscape at transcriptional resolution, we reanalyzed a publicly available scRNA-seq dataset (GSE164241; 13 GM and 8 PD subjects; 2 PD donors contributed multiple biopsies: PD164, three samples; PD161, two samples). C) UMAP visualization of nine transcriptionally distinct clusters identified among all T cells, and cell density in UMAP space, split by condition (GM vs. PD). D) Dot plot showing expression of curated marker genes (right) across the nine T cell clusters (C1–C9, bottom), grouped into seven functional categories and organized under CD8- and CD4-lineage headers; C8 (MAIT cells), though transcriptionally distinct from the conventional CD8 lineage, is grouped under the CD8 header for display purposes. Dot color represents scaled average expression per cluster; dot size represents the percentage of cells within each cluster expressing the gene. E) Donut plots showing the proportion of cells from each of the nine T cell clusters in GM and PD, expressed as a percentage of total T cells per condition. F) Volcano plot of subject-level pseudobulk differential expression analysis comparing PD vs. GM for all T cells. G) Differential abundance of T cell clusters between PD and GM, showing odds ratio and 95% confidence interval for each cluster. Clusters enriched in PD (OR > 1) are shown in red; clusters enriched in GM (OR < 1) are shown in green. Statistical significance was assessed at the sample level, patient level, and donor-blocked level using propeller with FDR correction (Supplementary table 1).

To characterize the transcriptomic landscape of antigen-experienced CD4⁺ T cells in GM and PD, we analyzed a publicly available scRNA-seq dataset comprising 13 GM and 8 PD samples from 18 donors (two donors contributed multiple biopsies: PD164, three samples; PD161, two samples; GSE164241 (25)). We isolated the T cell compartment by sequential unsupervised clustering of gingival immune cells followed by identification of T cells based on canonical marker expression, yielding nine transcriptionally distinct clusters (C1–C9) with similar representation across GM and PD (**Figure 1C–E**). Five clusters corresponded to the CD4⁺ lineage: C2 (naïve/central memory-like, the most frequent CD4^+^ cluster), C6 (activated/memory-like), C5 (Treg-like), C4 (Th17/Treg-like), and C9 (Tfh/Tph-like). Three clusters corresponded to the CD8⁺ lineage: C1 (effector/cytotoxic-like, the most frequent CD8⁺ cluster), C3 (resting/effector-memory-like), and C7 (terminally differentiated effector-like). C8 was identified as MAIT cells based on high expression of SLC4A10, ZBTB16, and KLRB1 (26).

Because cluster frequencies did not differ significantly between PD and GM (**Figure 1E**), we asked whether disease was instead reflected by a transcriptional state and therefore performed pseudobulk differential expression analysis (n= 21 [8 PD, 13 GM]; 10,796 genes after filtering) (**Figure 1F**). This identified 2,878 genes at FDR < 0.05 (616 with >2-fold change; 247 higher in PD, 369 higher in GM). In GM, the differentially expressed genes were broadly associated with naïve/resting lymphocyte cytoskeletal and adhesion programs (CD52, PFN1, JAML, MYL12A, LPAR6, CKLF, LGALS1, PSMA4, TMSB4X). In PD, the differentially expressed genes were downstream of TCR and cytokine signaling, consistent with chronic antigen-driven activation. Notably, several Tfh/Tph-associated genes (CXCR5, CXCL13, BCL6, TOX2) were detected in the bulk T cell signal in PD, despite arising from a minority subpopulation.

To determine whether the Tfh/Tph-like C9 cluster was specifically associated with PD, we performed a cluster abundance analysis and calculated odds ratios for each cluster in PD versus GM (**Figure 1G**). Of all clusters, C9 showed the largest PD-associated shift in abundance (from 2.5% to 4.9% of T cells, OR = 2.26; 95% CI: 1.27–4.01), reaching significance at the sample level (FDR= 0.026) and patient level (FDR= 0.046), with the association lessening to a non-significant association after donor-blocking, which accounts for repeat biopsies from the same donor (within-donor correlation ρ = 0.73; FDR= 0.064). All other clusters had donor-blocked FDR values between p= 0.5–0.9 (**Supplementary Table 2**). Together, the data suggests the disease-associated change was not a quantitative increase in the numbers of antigen-experienced cells, but rather a qualitative reprogramming within an already-resident pool.

### The Sensitized Recurrent Periodontitis model (SeRPD) promotes gingival T cell enrichment while preserving bone healing

Given the recurrent nature of human periodontitis, we asked whether temporary immune dysregulation contributes to long-term tissue retention of antigen-experienced T cells in tissue. To test this hypothesis, we developed a sensitized recurrent periodontitis model in mice (SeRPD). The model consisted of four phases: Baseline (BAS), Periodontitis (PD), Recovery (REC) and Recurrence (RCU). The phases were induced by bilateral placement, removal and replacement of silk ligatures around second maxillary molars. The PD phase lasted 10 days, followed by 30 days of recovery, and lastly 10 days of recurrence. This timeline allowed us to define a “recovered” reference state rather than a healthy baseline, an approach that is more clinically relevant. Transient Treg depletion was used as an experimental tool to enhance T cell priming during active disease-without inducing permanent immune dysregulation-by administering diphtheria toxin to Foxp3^DTR^ mice on days 1and 2 post-ligature placement, restricted to the PD phase (**Figure 2A**). The depletion was short-lived: compared with baseline and the maximum depletion day (day after DT application, MaxDep), we observed that at day 10 (end of PD phase) an average of 37.4% of the Tregs (TCRβ^+^CD4^+^Foxp3^+^CD25^+^) had repopulated the cervical lymph nodes (CLN), and by the end of the recovery phase (day 40), the Treg compartment was fully restored at CLN (**Figure 2B, C**). Gingival mRNA levels of Foxp3 were significantly higher than baseline after recovery (**Figure 2D**), indicating that there were no lingering regulatory deficits after recovery. Critically, the transient regulatory disruption did not compromise the alveolar bone recovery, as confirmed by micro-CT quantification (**Figure 2E**).

**Figure 2.**
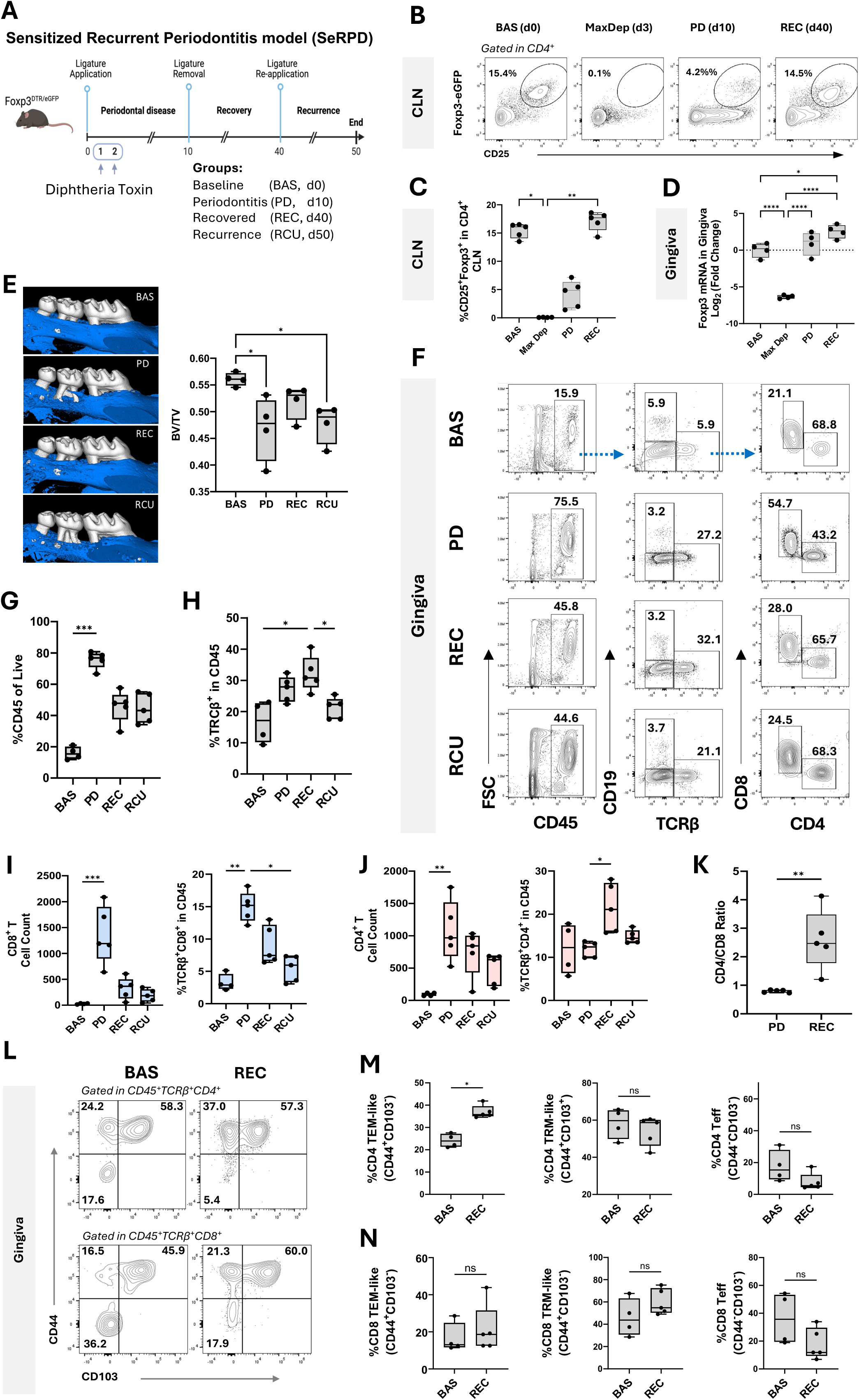
The Sensitized Recurrent Periodontitis model in mice (SeRPD) promotes gingival T cell enrichment while preserving bone recovery. A) Experimental design of the Sensitized Recurrent Periodontitis model in mice (SeRPD). B) Representative dot plots and mean percentage of Foxp3^+^CD25^+^ within CD4^+^ gate in cervical lymph nodes at Baseline (BAS, n=5), maximum Treg depletion (MaxDep, after two DT applications; n=4), Periodontitis (PD, day 10, n=5) and Recovery (REC, day 40; n=5). C) Frequency of Foxp3^+^CD25^+^ within CD4^+^ in cervical lymph nodes. D) Gingiva mRNA expression of *Foxp3* by RT-qPCR, normalized to β-Actin (n=5 per group). E) Representative microCT reconstructions of maxillary bone from each group (blue volume: alveolar bone) and quantification of bone volume (BV) to total volume (TV) ratio (n=4 per group). F) Representative flow cytometry gating strategy of four concatenated independent samples; numbers indicate mean frequency of each population within its parent gate. Dashed blue lines indicate gating sequence. G) Frequency of CD45⁺ cells in gingival tissue. H) Frequency of TCRβ⁺ cells in CD45⁺. I) Total count and frequency of live CD45⁺TCRβ⁺CD8⁺ T cells. J) Total count and frequency of live CD45⁺TCRβ⁺CD4⁺ T cells in gingiva. K) CD4⁺/CD8⁺ ratio during periodontitis (PD) and recovery (REC) phases. L) Representative density plots and mean percentage of CD44^+^, CD44^+^CD103^+^ and CD44^-^CD103^-^ within CD45^+^TCRβ^+^CD4 and CD45^+^TCRβ^+^CD8^+^ gates. M and N) Percentages of TEM-like (CD44^+^CD103^-^), TRM-like (CD44^+^CD103^+^) and Teff (CD44^-^CD103^-^) in CD4^+^ and CD8^+^ cells. Sample size in F-N: BAS n=4; PD n=5; REC n=5; and RCU n=5. Statistical comparisons were performed using the Mann–Whitney U test for pairwise comparisons and the Kruskal–Wallis test with post hoc correction for multiple comparisons. In all graphs, each dot represents one biological replicate (one mouse). ns = *p* > 0.05, * *p* < 0.05, ** *p* < 0.01.

Flow cytometry analysis of gingival tissue (**Figure 2F; Supplementary Figure 1**) revealed a pronounced immune cell infiltration (CD45⁺) that peaked during the PD phase (**Figure 2G**). After disease resolution, T cells (CD45⁺ TCRβ⁺) remained locally enriched in the gingiva (**Figure 2H**). Total T cell infiltration was markedly elevated at PD and predominantly composed of CD8⁺ T cells, consistent with transient regulatory disruption (**Figure 2I, J**). After recovery, both CD8^+^ and CD4⁺ T cells remained at significantly higher levels than baseline. To evaluate differential relative T cell retention, we calculated the CD4/CD8 T cell ratio, normalizing for inter-individual variability in abundance. This ratio increased substantially after recovery, rising from a median of 0.8 during PD to 2.5, consistent with a preferential shift toward CD4^+^ T cell enrichment after recovery (**Figure 2K**). To assess whether gingivally retained T cells display a memory-like phenotype after recovery, we compared the frequencies of CD4^+^ and CD8^+^ effector-memory-like (TEM-like) and resident-memory-like (TRM-like) cells within their respective compartments (CD45^+^TCRβ^+^CD4^+^/CD8^+^) in recovered versus baseline (**Figure 2 L-N**). Only CD4^+^ CD44^+^CD103^-^ TEM-like frequency was significantly increased in recovered mice (**Figure 2M**). Together, these findings indicated that sensitized periodontitis was followed by a lasting enrichment of antigen-experienced, memory-phenotype CD4^+^ T cells in gingiva after structural and regulatory recovery, consistent with immune remodeling by prior inflammatory exposure.

To contextualize the T cell findings within the broader gingival immune landscape, we also analyzed the myeloid compartment (CD45^+^CD19^-^TCRβ^-^CD3^-^TCRγδ^-^) (**Supplementary Figure 1**). We observed higher frequencies of neutrophils (LY6G^+^CD11b^+^) and activated monocytes (CD11b^+^LY6C^+^MHCII^+^) in recurrence, particularly compared with recovery and PD phases. In contrast, both the recovered and baseline groups displayed higher proportions of putative macrophages (CD11b^+^CD11c^-^LY6G^-^LY6C^-^MHCII^var^). Notably, conventional dendritic cells (cDCs; CD11c^+^MHCII^+^) were markedly increased in the recovered group and, across all conditions, were mostly cDC2 subset (CD103^-^). These changes provide contextual information on immune and tissue remodeling across phases. In contrast to T cells, B cells (CD45^+^CD19^+^) did not expand during PD or remain retained in the gingiva after recovery, and their relative frequency within the flow-defined CD45^+^ compartment remaining stable across experimental groups (**Supplementary Figure 2**), suggesting the model preferentially imprinted T cell-dominant rather than humoral immune changes.

### Transient regulatory breakdown increases regional CD4⁺ TEM frequency and enhances mucosal T cell retention, but periodontitis history alone is sufficient to increase TEM pathogenicity

To determine how prior regulatory disruption shapes outcomes after recovery, we compared two groups of recovered animals: those with a history of Treg depletion during PD (REC) and those without (Recovery non-depleted: RND), against baseline controls. Morphometric analysis showed no difference in alveolar bone recovery among REC, RND, and baseline mice (**Figure 3A**). In the gingiva, T cell retention after recovery increased with previous regulatory breakdown (**Figure 3B, C**). To test whether this inflammatory history also shaped regional memory T cell formation, we compared memory subsets frequencies in cervical lymph nodes (CLN), of REC and RND mice against their respective baseline littermate controls. In RND, CD4^+^ effector memory T cell (TEM) frequency was significantly increased and CD4^+^ naïve T cell frequency significantly decreased relative to controls (**Figure 3D, E**); CD8^+^ TEM frequency was not significantly altered, while CD8^+^ central memory T cell (TCM) frequency decreased (**Figure 3F**). In REC, CD4^+^ TEM, CD8^+^TEM, and CD8^+^ TCM frequencies were all significantly increased, and CD4^+^ naïve frequency decreased, relative to controls (**Figure 3G-I**). To directly compare the magnitude of RND and REC responses, we calculated the Log_2_ fold-change relative to baseline for each memory subset within the CD4 and CD8 lineages. Thus, prior regulatory disruption significantly amplified memory subset formation and decreased naïve T cell frequency in REC relative to RND, except for CD4^+^ TCM which was unchanged (**Figure 3J, K**).

**Figure 3.**
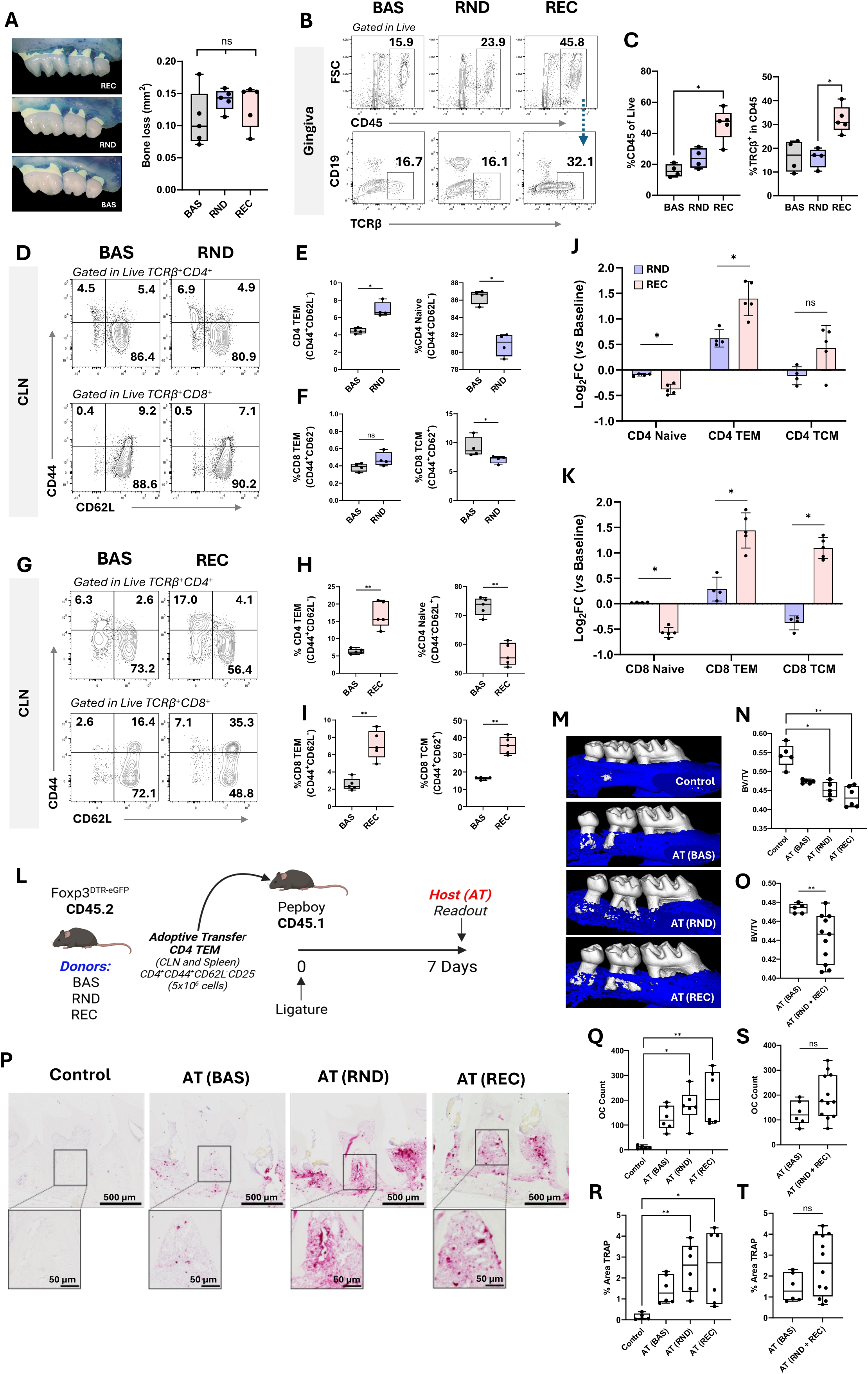
Transient regulatory breakdown increases regional CD4⁺ TEM frequency and enhances mucosal T cell retention, but periodontitis history alone is sufficient to increase TEM pathogenicity. A) Morphometric analysis of alveolar bone volume in baseline (BAS) mice and in recovery groups that underwent ligature-induced periodontitis either with transient DT-mediated Treg depletion (REC) or without depletion (RND); n=5 per group. B) Representative dot plots of four concatenated gingival samples per group (BAS, RND, REC) showing leukocytes (CD45⁺) and T cells (CD45⁺TCRβ⁺). The dashed blue arrow indicates the gating sequence; numbers indicate the mean frequency of each population within its parent gate. C) Frequency of CD45⁺ cells within the live gate and of TCRβ⁺ cells within CD45⁺ cells (BAS n=4, RND n=4, REC n=5). D) Representative flow cytometry plots of cervical lymph node (CLN) samples from the BAS (n=4) and RND (n=4) groups; numbers indicate the mean frequency of each population within its parent gate, as indicated by the italicized labels (live TCRβ⁺CD4⁺ or live TCRβ⁺CD8⁺). E) Frequency of CD4⁺ effector memory (TEM: CD44⁺CD62L⁻) and naïve (CD44⁻CD62L⁺) CD4⁺ T cells in CLN. F) Frequency of CD8⁺ TEM (CD44⁺CD62L⁻) and CD8⁺ central memory (TCM: CD44⁺CD62L⁺) cells in CLN. G–I) As in D–F), comparing the REC (n=5) group with its respective baseline (n=5). J, K) Log₂ fold-change in CD4⁺ J) and CD8⁺ K) CLN subpopulations in the RND and REC groups relative to their respective experimental baselines. L) Experimental design of the adoptive transfer experiment. M) Representative µCT images of each host group 7 days after transfer: control (untouched; no ligature, no transfer), AT (BAS), AT (RND) and AT (REC), indicating transfer of CD4⁺ TEM cells isolated from BAS, RND and REC donors, respectively. N) Bone volume/total volume (BV/TV) ratio across all groups. O) BV/TV ratio comparing AT (BAS) with pooled AT (RND) + AT (REC) groups. P) Representative TRAP staining with corresponding insets showing osteoclasts around the furcation of the second molar in each experimental group. Q, R) Number of osteoclasts within the region of interest across all groups Q) and in AT (BAS) versus pooled AT (RND) + AT (REC) S). R, T) TRAP⁺ stained area (% of region of interest) across all groups R) and in AT (BAS) versus pooled AT (RND) + AT (REC) T). Statistical comparisons were performed using the Mann–Whitney U test for pairwise comparisons and the Kruskal–Wallis test with post hoc correction for multiple comparisons. In all graphs, each dot represents one biological replicate (one mouse). ns = *p* > 0.05, * *p* < 0.05, ** *p* < 0.01.

Since the CD4^+^ TEM subset was significantly increased in both REC and RND, we asked whether these cells differed in pathogenicity. We transferred 5×10^6^ TEM cells (defined as CD4^+^CD44^+^CD62L^-^CD25^-^) isolated from CLN and Spleen of REC, RND or baseline donor mice (Foxp3^DTR^, CD45.2) retro-orbitally into congenic CD45.1 mice, followed by 7 days of ligature placement; untouched mice with no ligature and no cell transfer served as controls (**Figure 3L**). Both micro-CT analysis of the bone loss (**Figure 3M-N**) and TRAP analysis of osteoclasts around the second molar (**Figure 3P, Q, R**) showed that recipients of REC- or RND-derived TEM cells (AT REC, AT RND) displayed significantly higher bone loss and osteoclast numbers detection compared to untouched controls, with no significant difference between the two groups.

To test the broader hypothesis that any prior periodontitis exposure shapes TEM pathogenicity, we compared AT(BAS) against the pooled RND- and REC-derived TEM recipients (AT(RND+REC), n=11). This pooled comparison revealed significantly greater bone loss in AT(RND+REC) recipients than AT(BAS) (p = 0.005), suggesting that periodontitis history, rather than regulatory breakdown, primarily drives TEM pathogenicity in this model (**Figure 3O**). This difference was not accompanied by a corresponding significant increase in osteoclast numbers at the tissue level (**Figure 3S, T**), suggesting that reduced bone volume in recipients of disease-experienced donor TEM cells may reflect cumulative osteoclast activity over the course of the experiment instead of a difference detectable at day 7 post-adoptive transfer.

### High-resolution T cell profiling through paired single-cell transcriptomics and TCR sequencing confirms gingival enrichment of CD4^+^ T cells upon recovery

To analyze the transcriptomic landscape of the gingival T cells in the SeRPD model, we performed paired 5′ scRNA-seq with TCR amplification. (**Figure 4A**). Given the experiment design, proportions reflect relative immune composition within the CD45⁺ compartment. Based canonical and population-restricted gene expression, we identified 4 major cell classes (T cells, B cells, Myeloid cells and Stromal (likely contamination)) (**Figure 4B and Supplementary Figure 2A-B**). Consistent with the flow cytometry data, both PD and recovered groups displayed the highest proportion of T cells in the gingiva (PD: 41.9% and REC: 45.2%). The highest B-cells proportion was detected in baseline (19%), while the myeloid compartment was the largest in recurrence (66.8%), signifying a relative compositional shrinkage as T and myeloid compartments expand during disease, rather than a true change in B-cell numbers (**Figure 4B**). Further analysis of the myeloid compartment revealed a distinct population of type I interferon (IFN-I)–stimulated neutrophils that became dominant during the PD phase (55%), occurring in the context of the transient Treg depletion (**Supplementary Figure 3**). During recovery, the immune proportions returned toward homeostatic distribution, resembling baseline. Finally, in the recurrence phase, macrophages/monocytes and neutrophil heterogeneity re-emerged (**Supplementary Figure 3**).

**Figure 4.**
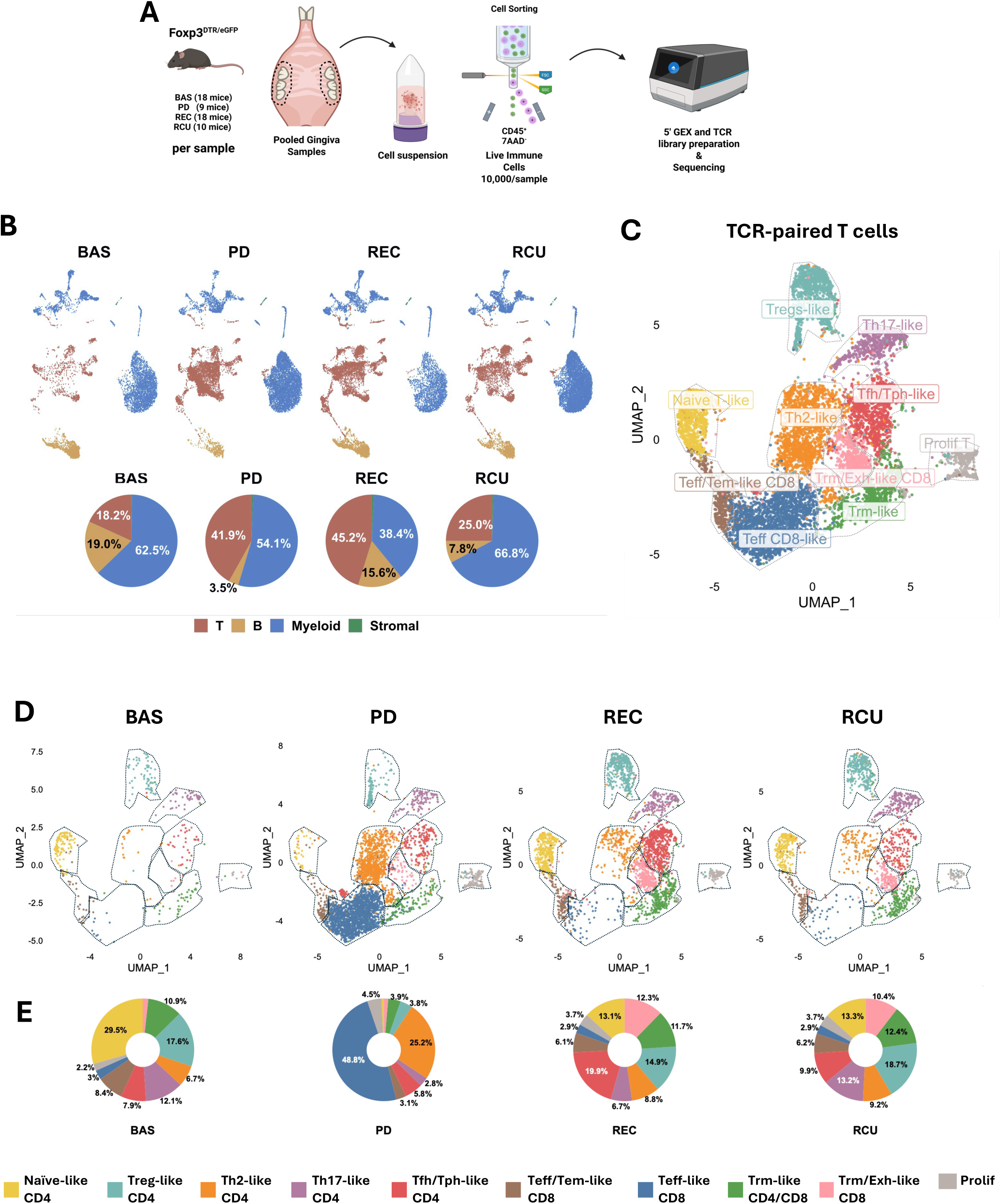
High-resolution T cell profiling through paired single-cell transcriptomics and TCR sequencing confirms gingival enrichment of CD4^+^ T cells upon recovery. A) Experimental design. Gingival tissue was pooled to obtain ≥10,000 live CD45⁺ immune cells per sample (BAS n = 18, PD n = 9, REC n = 9, RCU n = 18 mice per pool), and two independent pools per condition were processed for 5′ gene expression and V(D)J sequencing. B) UMAP of 51,572 gingival cells split by condition (BAS: 8,142; PD: 14,072; REC: 13,349; RCU: 16,009), annotated into T cells, B cells, myeloid cells and stromal contaminants; pie charts show the proportion of each population per condition. C) UMAP of 9,224 αβ TCR⁺ T cells (both single and double TCR paired cells were included), after removal of non-T cell contaminants and unpaired T cells; annotated by gene-expression signatures into ten subclusters as (1) Naïve-like CD4 (naïve CD4 T cells); (2) Treg-like CD4 (regulatory T cells); (3) Th2-like CD4 (T helper 2 cells); (4) Th17-like CD4 (T helper 17 cells); (5) Tfh/Tph-like CD4 (T follicular helper/T peripheral helper–like cells); (6) Teff/Tem-like CD8 (effector/effector memory–like CD8 T cells); (7) Teff-like CD8 (activated effector CD8 T cells); (8) Trm-like CD4/CD8 (tissue-resident memory–like cells of mixed CD4/CD8 lineage); (9) Trm/Exh-like CD8 (tissue-resident memory/exhausted-like CD8 T cells); (10) Prolif (highly proliferative T cells). D) The UMAP in (C) split by condition (BAS: 404 cells; PD: 3,958; REC: 3,042; RCU: 1,820). Subcluster boundaries are marked by black outlines. E) Pie charts showing the proportion of each T cell subcluster within each condition, as a percentage of all αβ TCR⁺ T cells in that condition.

Having established the broader immune gingival landscape, we focused on the T cell cluster-after removing NK cells and innate contaminants-paired with their corresponding TCRα and/or β chain sequences. Across groups, an average of 42% of T cells lacked a detectable TCR chain, with the baseline group exhibiting the highest proportion at 60.4% (**Supplementary Figure 2C**). TCR dropout correlated with sequencing depth within each condition, but sequencing depth alone did not explain the elevated dropout rate in baseline, since BAS T cells showed comparable depth distributions to REC and RCU (**Supplementary Figure 2D**). This finding suggested a depth-independent condition effect, potentially linked to the smaller T cell fraction at baseline (18.2%).

Unbiased sub-clustering of TCR-paired T cells revealed ten transcriptionally distinct populations (**Figure 4C**): 1) naïve-like CD4⁺ T cells; 2) regulatory T cell (Treg)-like CD4⁺; 3) T helper type 2 (Th2)-like CD4⁺; 4) T helper type 17 (Th17)-like CD4⁺; 5) T follicular helper/T peripheral helper (Tfh/Tph)-like CD4⁺; 6) effector/effector memory T cell (Teff/Tem)-like CD8⁺; 7) effector T cell (Teff)-like CD8⁺; 8) tissue-resident memory T cell (Trm)-like CD4⁺/CD8⁺; 9) tissue-resident memory/exhausted T cell (Trm/Exh)-like CD8⁺; and 10) proliferating cells. The annotation was established by the combined expression of phenotypically distinctive genes as indicated in **Supplementary Figure 4**. The Tfh/Tph-like annotation was assigned based on a co-occurring gene module including *Cxcr6*, *Bcl6*, *Il21*, *Icos*, *Cd40lg, Batf,* and *Pdcd1* (**Supplementary Figure 4A, B**), indicating that the data do not clearly separate follicular vs. extrafollicular helper cells at this resolution. While Cxcr5 expression was negligible in this cluster, a subset retained *Bcl6* expression (17%), the lineage-defining transcription factor for Tfh differentiation (**Supplementary Figure 4C**). We therefore favored a Tph-skewed annotation for this cluster, rather than an equal split between the two subtypes. The Trm-like CD4/CD8 mixed-lineage population was annotated based on a shared Trm gene core and contains 66% of CD8^+^ cells and 34% of CD4^+^ cell. This lineage composition was not resolved by unbiased clustering, which instead split the cluster based on activation score rather than lineage; thus, we chose to preserve it as a single mixed cluster (**Supplementary Figure 4D**).

When we evaluated these populations across conditions (**Figure 4D, E**), we observed that, at baseline, the largest groups were Naïve-like CD4⁺ cells (29.5%) and Treg-like (17.6%). During PD, as previously shown by flow cytometry, effector CD8⁺ T cells were markedly predominant (48.8%), accompanied by expansion of Th2 CD4⁺ T cells (25.2%). Following recovery (REC), the T cell landscape partially returned toward baseline composition, though it remained distinct from baseline. The CD4⁺ compartment was enriched for Tfh/Tph-like cells (19.9%), Treg-like (14.9%), and Naïve CD4⁺ cells (13.1%), along with Trm-like CD4/CD8 cells (11.7%). At recurrence (RCU), the T cell profile shifted slightly relative to recovery, with increased proportions of Th17-like (13.2%), and Treg-like (18.7%) cells, indicating immune reactivation skewed toward a Th17 transcriptomic profile.

### Gingivally retained clonally expanded CD4^+^ T cells exhibit a Tfh/Tph-like transcriptomic signature and persist through recurrence

We further refined our T cell analysis to account for clonal selection of gingiva-retained T cells; for this, we mapped clonally expanded T cells across conditions using a stringent criteria. Expanded T cells were defined as 2 or more different cell barcodes with identical TCRα and β gene usage and CDR3 (hypervariable segment within the TCR that directly binds to epitope) nucleotide sequences per sample. Unique TCR α and βgene usage and/or different CDR3 nucleotide sequences per sample were considered non-expanded (**Figure 5A**). We detected that 8.2% of all TCR-paired T cells in BAS were clonally expanded, 16.5% in PD, 20.6% in REC, and 19.3% in RCU (**Figure 5B**). We then mapped these expanded T cells onto the previously defined T cell clusters and identified the dominant populations within each group (**Figure 5C**). In BAS, the few clonally expanded T cells detected were mostly tissue-resident CD8^+^ T cells, with Trm/Exh-like CD8 T cells showing the highest expansion frequency (66.7% of the cluster), followed by Trm-like cells (21.2%; **Figure 5D**). In PD, most of the expanded T cells were effector CD8^+^, representing 56.6% of all expanded cells. Only 19.4% of the Teff CD8-like cluster itself was clonally expanded (**Figure 5E**); indicating massive infiltration of largely non-expanded (newly recruited) effector CD8^+^ T cells during the temporary Treg disruption. These expanded CD8^+^ T cells did not persist in the tissue through recovery, where most expanded T cells were CD4 ^+^ Tfh/Tph-like cells (42.2% of all expanded T cells); notably, 44.9% of the Tfh/Tph-like cluster itself was expanded at this timepoint. A substantial proportion of these Tfh/Tph-like cells persisted through recurrence, where 32.6% of the cluster remained clonally expanded. At recurrence, other clusters also showed elevated within-cluster expansion, including Th17-like (23.6%) and CD8 Trm populations (Trm-like: 27.9%; Trm/Exh-like CD8: 39.1%; **Figure 5D, E**). These results indicated that each phase was characterized by distinct proportions of expanded T cell subsets, with both the recovery and recurrence phases showing higher CD4^+^ T cell clonal expansion.

**Figure 5.**
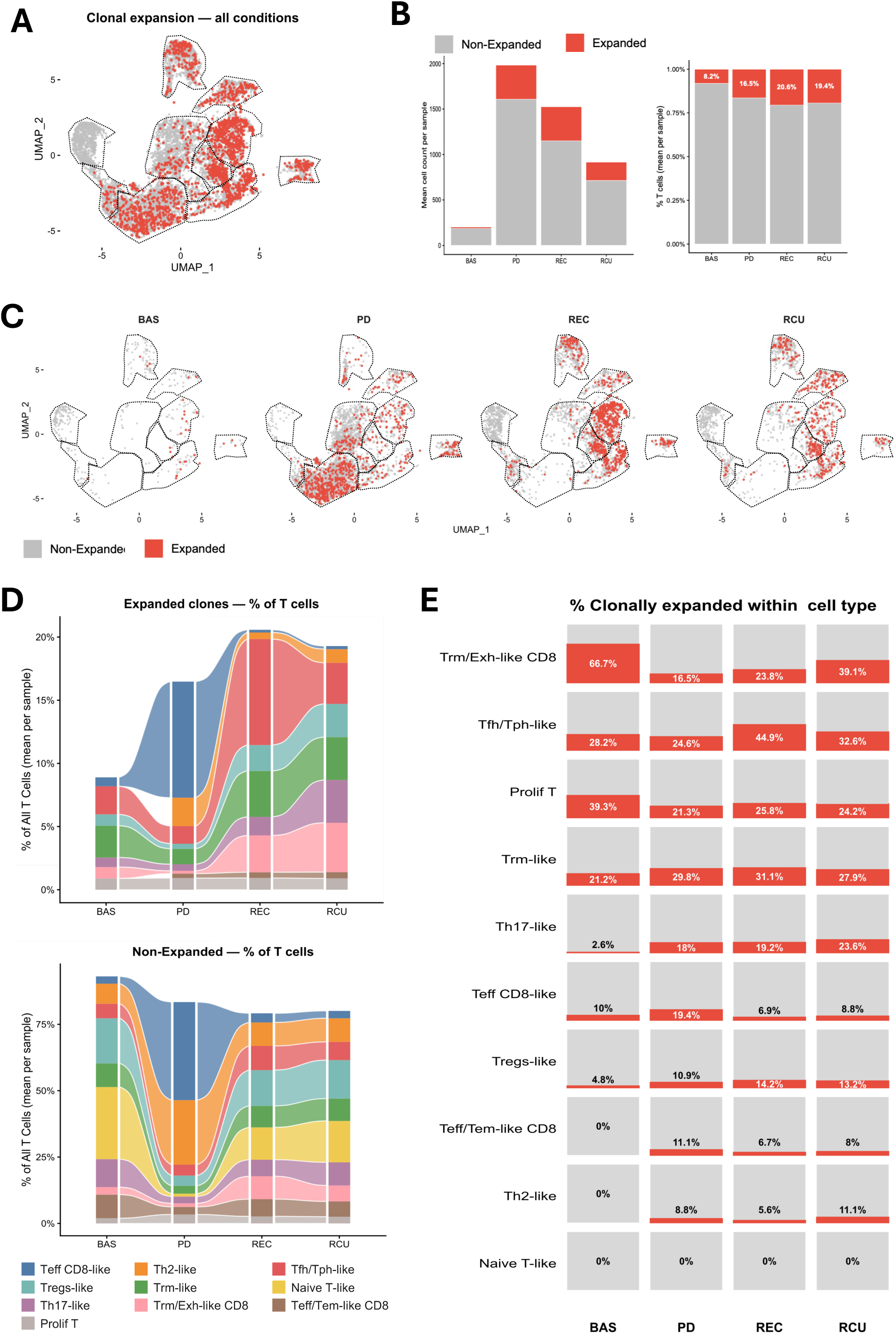
Gingivally retained clonally expanded CD4^+^ T cells exhibit a Tfh/Tph-like transcriptomic signature. A) UMAP of αβ TCR⁺ T cells (n = 9,224) from all conditions, highlighting clonally expanded (red) and non-expanded (grey) cells; outlines indicate the T cell subclusters defined in Figure 4C. B) Mean cell counts (left) and mean percentages (right) of expanded and non-expanded T cells per sample in each condition; numbers indicate the percentage of expanded cells. C) Condition-stratified UMAPs showing the distribution of expanded cells across T cell subclusters. D) Alluvial plots showing the subcluster composition of expanded (top) and non-expanded (bottom) T cells, expressed as a percentage of all αβ TCR⁺ T cells per sample. E) Percentage of clonally expanded cells within each T cell subcluster per condition. Expanded T cells were defined as ≥2 cells within the same sample sharing an identical clonotype (identical TCR gene usage and CDR3 nucleotide sequence).

Given the continued presence of expanded CD4^+^ T cells clusters, we asked whether these cells conserved their TCR architecture through these phases. The mouse TCR is produced by somatically recombining gene segments named variable (V), junctional (J), diversity (D) and constant (C) that assemble into alpha and beta chains (**Figure 6A**). First, we evaluated V (TRBV) and J (TRBJ) gene pairing in the beta chains from expanded and non-expanded CD4^+^T cells across disease phases (**Figure 6B**). We confirmed that as the disease progresses and more clones expand, the expanded compartment retains a biased subset of V-J pairings relative to the full repertoire (**Figure 6B**). In particular, some of these pairings (e.g., TRBV5-TRBJ1-4 combination) reappear in REC and RCU. To quantify beta-chain similarity between recovery and recurrence, we identified each unique beta-chain gene set and compared their prevalence in both groups, identifying genes enriched in REC and RCU, and 6 unique beta chains with similar prevalence in both phases (**Figure 6C**). To measure how many beta chains present at recovery reappear at recurrence per CD4 cluster (**Supplementary Table 3**), we calculated the Jaccard index as a repertoire-convergence metric (**Figure 6D**). The global Jaccard index across all CD4^+^ T cells was 0.2, indicating that 20% of the combined (union) set of clonotypes was shared between the two timepoints. The Tfh/Tph-like cluster had the highest Jaccard index amongst individual CD4 clusters (0.132), indicating that 13% of its beta chains recur at both timepoints, rather than the cluster being freshly repopulated from a naïve pool at each phase (**Figure 6D**). Taken together, the TCRβ repertoire analysis revealed a subset of highly expanded clonotypes that were shared across recovery and recurrence cohorts. Because these disease phases were analyzed in distinct animals, these shared clonotypes represent public, expanded T cell populations rather than longitudinally persistent clones.

**Figure 6.**
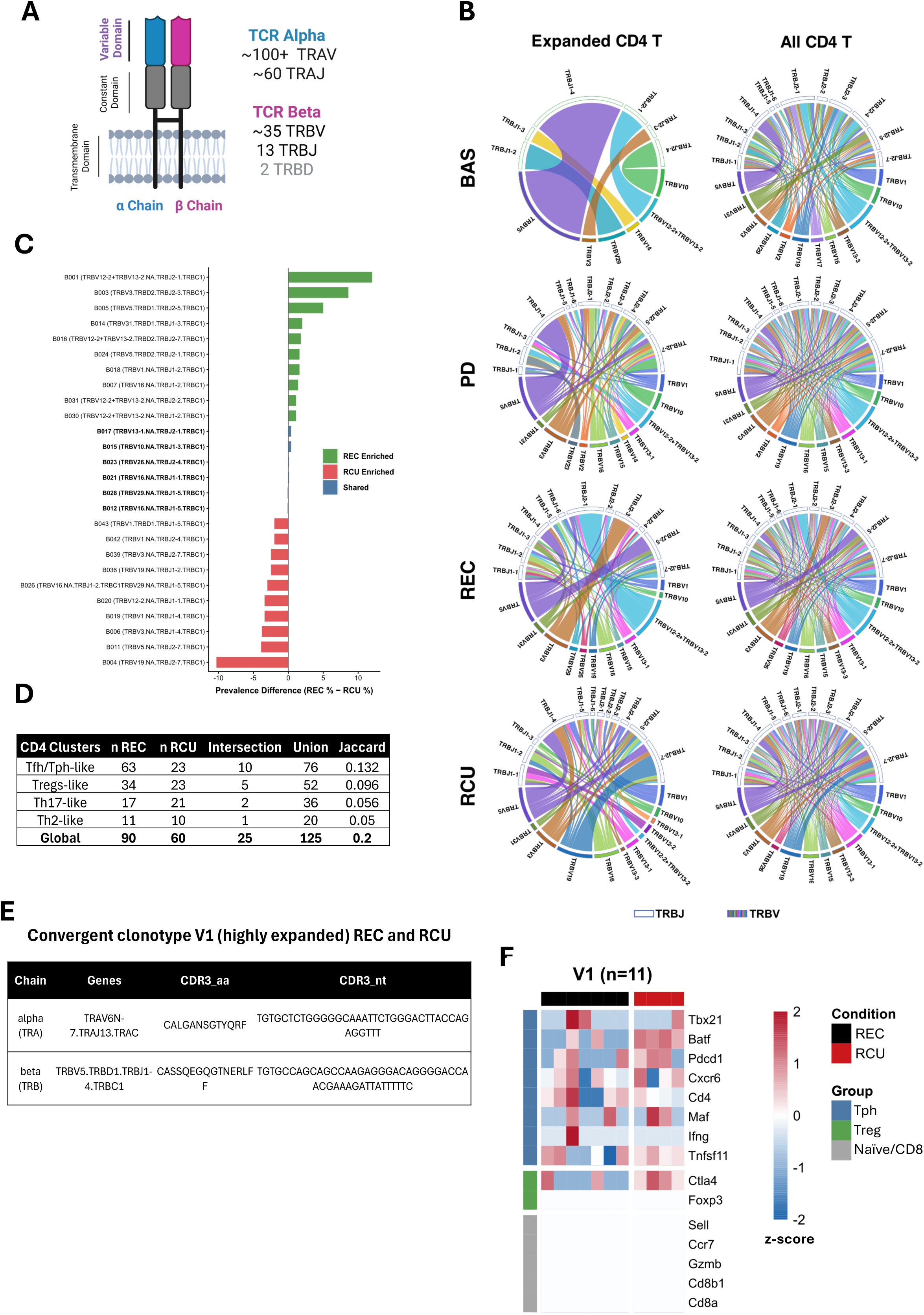
Gingival clonally expanded Tfh/Tph-like CD4^+^ T cells clones converge in recovery and recurrence. A) Schematic of the αβ TCR showing the variable and constant domains of the α and β chains and the approximate number of mouse germline V, D and J gene segments. B) Chord diagrams of TCRβ V–J gene pairing in expanded (left) and all (right) CD4⁺ T cells in each condition. In each diagram, colored arcs (bottom) represent TRBV genes and white arcs (top) represent TRBJ genes; ribbon width is proportional to the frequency of each V–J pair within the condition. The 12 most frequent TRBV and TRBJ genes are shown per diagram, and only cells with a detected TCRβ chain are included. CD4⁺ T cells comprise the Tfh/Tph-like, Treg-like, Th17-like and Th2-like subclusters. C) Difference in the prevalence (REC % − RCU %) of TCRβ gene combinations (TRBV–TRBD–TRBJ–TRBC) among expanded CD4⁺ T cells in REC (n = 442) and RCU (n = 203). Each unique combination was assigned a code (B001, B002,…) in order of overall prevalence (Supplementary table 2). Shown are the ten most REC-enriched (green) and RCU-enriched (red) combinations and all shared combinations (blue). Combinations were classified as enriched when the prevalence difference exceeded 0.5 percentage points, and as shared when the difference was ≤ 0.5 percentage points and the combination was detected in ≥ 3 cells in either condition. D) Jaccard similarity of TCRβ gene combinations between REC and RCU expanded CD4⁺ T cells, per CD4 subcluster and for all CD4⁺ T cells combined (Global), with the number of distinct combinations in each condition, their intersection and their union. E) TCRα and TCRβ gene usage and CDR3 amino acid and nucleotide sequences of clonotype V1, which was detected with an identical nucleotide sequence in both REC and RCU. Expanded T cells are defined as in Figure 5 (Supplementary Tables 3 and 4). F) Expression of selected Tph-associated, Treg-associated and negative marker genes in individual REC (n = 7) and RCU (n = 4) cells carrying clonotype V1; expression is z-scored per gene across cells.

To further test the clonotype convergence, we evaluated identical clonotypes (alpha and beta chains) in both REC and RCU phases and found a highly expanded clonotype (V1, n=11 cells). We tracked the paired alpha/beta TCR sequence defining this clone. Confirmation of the paired amino acid and nucleotide sequences of the CDR3 region indicated that the convergence occurred at the DNA rearrangement level, not merely a coincidental amino-acid match from different underlying recombination events (**Figure 6E**). This clonotype predominantly belonged to the Tfh/Tph-like cluster and expressed the module *Tbx21*, *Batf*, *Pdcd1*, *Cxcr6*, *Cd4*. Notably, in recurrence, this clonotype also expressed Tnfsf11, suggesting participation in bone destruction (**Figure 6F**). Two additional expanded clonotypes (V2 and V3) were detected; both of which differ from V1 by a single nucleotide in the CDR3α or β sequences but encode an identical αβTCR protein (**Supplementary Tables 3 and 4**). The V2 variant was also detected at baseline, indicating that these clones may arise from pre-existing T cells within the normal gingiva. Across the combined 47 cells carrying this TCR, 51.1% mapped to the Tfh/Tph-like cluster, 23.4% to Trm-like CD8/CD4, 12.8% to Th2-like, 8.5% to Treg-like, 2.1% to Teff CD8-like and 2.1% to proliferating T cells (**Supplementary Table 4**). The repeated detection of identical αβ clonotypes across independent pools of mice in recovery and recurrence, suggested that this expanded T cell population arose through convergent selection rather than stochastic recruitment.

### Human periodontitis-associated T cells and gingivally retained T cells in mice share a B-helper transcriptomic signature

After identifying the Tfh/Tph-like cluster as the one that clonally converged the most between recovery and recurrence in our model, we next asked whether this mouse T cell cluster was transcriptomically related to the human C9 T cell cluster, which showed a PD-associated shift in abundance (**Supplementary table 2**). To address this, we assessed the expression of a Tfh/Tph “B-cell help” gene module across both human and mouse T cell clusters. The module comprises 11 genes conserved between species, spanning: lineage and phenotype markers (CD4, CD200), costimulatory/synapse machinery (ICOS, CD40LG, TNFRSF4, SH2D1A/SAP), the inhibitory receptor PD-1 (PDCD1), transcription factors (TOX2, BATF, MAF), and the effector cytokine IL21.

Beyond this conserved core, we included additional genes for each species based on detection threshold in the respective datasets. For humans, we included CXCL13 (B-cell chemoattractant), PASK (Tph-associated nutrient-sensing kinase), AFF3 (Tfh-associated in autoimmune studies), IL6ST (signaling upstream of STAT3, required to induce Bcl6/IL-21). In mice, we included *Bcl6* (Tfh master regulator), *Cd84* (Tfh-associated adhesion gene functionally linked to SAP), *Il21r* (mediates autocrine IL-21 signaling) and *Ebi3* (p28 partner subunit for IL-27, upstream of Bcl6/Tfh induction; **Figure 7A**).

**Figure 7.**
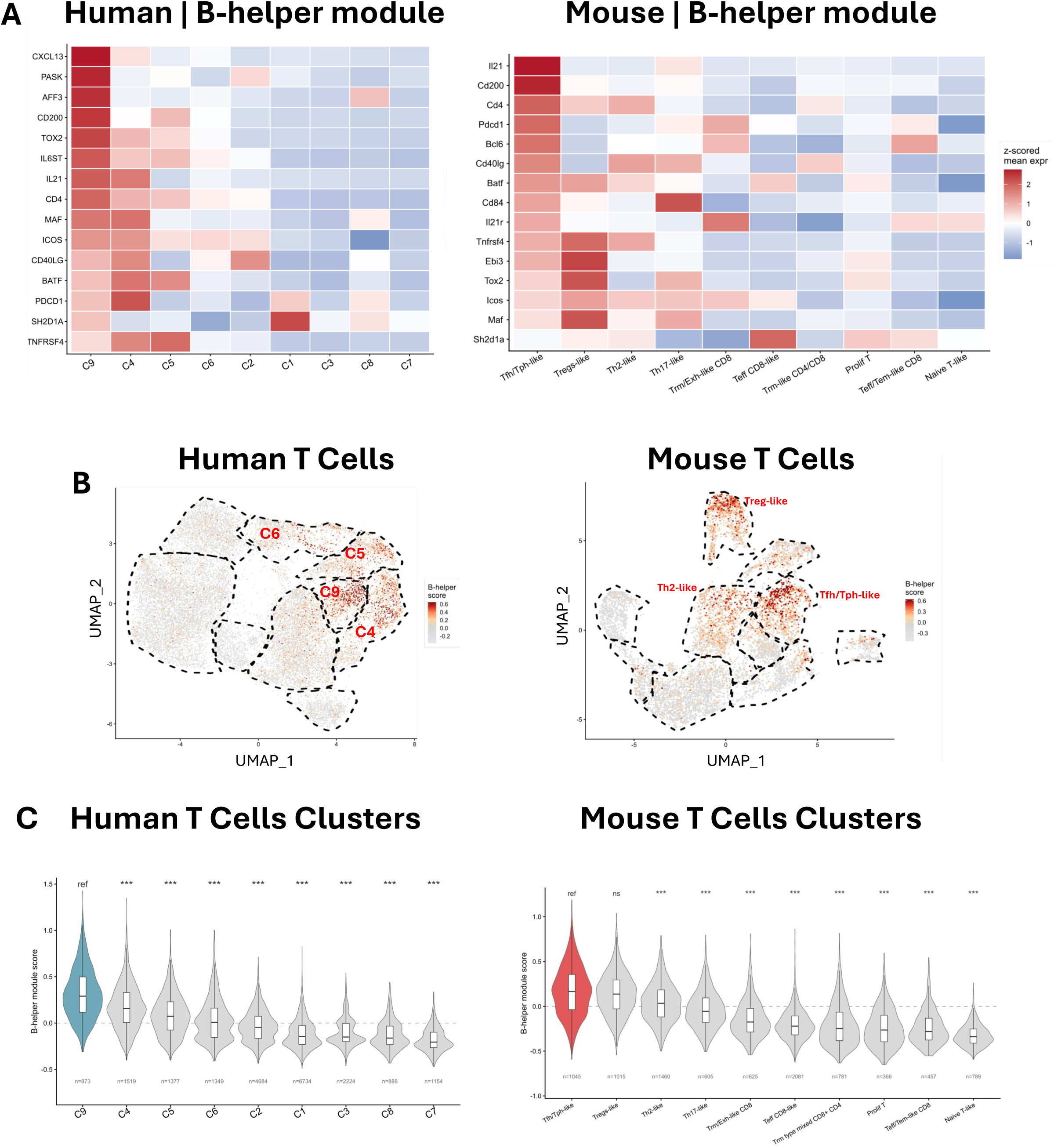
Human periodontitis-associated T cells and gingivally retained T cells in mice share a B-helper transcriptomic signature. A) Heatmaps of the 15 B-helper module genes across T cell clusters in human gingiva (left; clusters C1–C9 from Figure 1) and in the mouse SeRPD model (right; subclusters from Figure 4). Fill shows mean expression per cluster, z-scored within each gene. Clusters are ordered from highest (left) to lowest B-helper module score, and the target cluster (human C9; mouse Tfh/Tph-like) is outlined. B) UMAPs of human (n = 20,802) and mouse (n = 9,224) T cells colored by B-helper module score. The color scale is clipped to the 1st–99th percentile. Dashed lines outline clusters; the highest-scoring clusters are labeled. C) Distribution of B-helper module scores per cluster in human (left) and mouse (right), ordered by median score. Box plots show median and interquartile range, and the number of cells per cluster is shown below each violin. Module scores were calculated with Seurat AddModuleScore using species-specific 15-gene lists: 11 genes are shared, and each species has 4 genes of its own (see Methods). Scores are therefore compared only within each species. Statistical comparisons used the Kruskal–Wallis test followed by Dunn’s test with Benjamini–Hochberg correction, comparing each cluster with the target cluster (ref). *** p < 0.001; ns = not significant.

We confirmed that both clusters, C9 in human and the Tfh/Tph-like cluster in mouse, expressed the B-helper module (Figure 7A). We next compared module expression across the other T cell clusters by mapping the score onto each species’ respective T cell UMAP and observed that other clusters (C4 and C5 in human; Treg-like in mouse) also showed elevated scores ( **Figure 7B**). The C9 score was significantly higher than every other T cell cluster (p<0.001 for all comparisons), and in mouse the Tfh/Tph-like cluster was significantly higher than that of every other T cell cluster except the Treg-like cluster (p<0.001 for all comparisons except Treg-like, which was non-significant). This overlap with the Treg-like cluster was driven by high expression of genes also expected in Tregs (*Tnfrsf4, Icos, Ebi3, and Maf*) (**Figure 7C**). Thus, both clusters were robustly associated with a mixed Tfh/Tph phenotype, cell of this type function as master organizers of the B cell response in the tissue via IL-21 production and surface interactions.

## Discussion

In this paper, we show that transient immune regulatory breakdown during experimental periodontitis, followed by complete restoration of the regulatory compartment and structural recovery, leaves a lasting mucosal immune imprint: a clonally expanded CD4^+^ T population that reappears during a secondary inflammatory episode (recurrence). Because Treg restoration is complete and bone regeneration is preserved, long-term immune homeostasis is reestablished, supporting the interpretation that T cell enrichment reflects imprinting during the earlier window of regulatory disruption. Nucleotide-level convergence of TCRα and β CDR3 sequences across independent pools of mice indicates that this CD4^+^ T cell clonotype arises through convergent selection. The transcriptomic signature of highly expanded convergent clonotype also included the expression of Tnfsf11 (RANKL) during recurrence, suggesting a role in bone loss, with an overall phenotype consistent with the Tfh/Tph-like cluster. Together, these public clonotypes likely represent antigen-experienced, tissue-adapted T cells that are potentially recruited or expanded during recurrence.

Besides the Tfh/Tph reprogramming, our data showed that the sensitized model significantly increased the frequencies of regional effector memory CD4⁺ T cells and mucosa-enriched CD4⁺ T cells. Our adoptive transfer experiments indicated that this regulatory disruption is not required for these memory T cells to be pathogenic. Transfer of periodontitis-induced CD4⁺ memory T cells from secondary lymphoid organs increased bone loss in recipients regardless of the intensity of the initial inflammatory episode in the donor, suggesting that pathogenicity is an intrinsic property of periodontitis-induced T cell memory (24, 27, 28). Consequently, transient regulatory breakdown affects the recruitment and accumulation of an already-pathogenic population rather than its pathogenic potential. Thus, a history of regulatory disruption may promote recurrence by enlarging the pool of pathogenic memory T cells that persist after recovery.

Clonal convergence characterizes pathologies with sustained antigen exposure, including chronic infection, autoimmunity, atherosclerosis, and cancer, where persistent antigens drive the selection and convergent expansion of T cell clones (29–33). Periodontitis shares these features. Its pathogenesis is driven by reciprocal feedback between a dysbiotic microbiome and dysregulated inflammation, and in susceptible individuals the host response is unsuccessful and tissue-destructive (34, 35). The disease is also intrinsically persistent: current standard of care targets the biofilm, which rebuilds without regular maintenance; thus, a periodontitis patient is effectively a patient for life (10). Antigenic exposure is therefore sustained over decades. The T cell convergence at nucleotide level is consistent with shared antigen specificity, yet in our model the identity of the driving antigen remains unknown. Potential candidates include conserved epitopes from microbial components of the subgingival biofilm, self-antigens exposed during tissue destruction, or cross-reactive epitopes shared between the two (36).

Chronic inflammation may be accompanied by intermittent breaks in immune regulation. Several factors that increase periodontitis susceptibility, notably psychological stress and the hormonal shifts of pregnancy (37, 38), have recently been associated with transient Tregs redistribution or malfunction (39–41). Treg numbers fluctuate across the lifespan, and mucosal tissues are particularly sensitive to temporary declines in regulatory responses (2, 42). Thus, chronic antigenic exposure from the subgingival biofilm and intermittent transient regulatory disruptions from common life events can cumulatively enrich clonally convergent T cells in gingival tissue. Consistent with this, early studies showed that gingival T cell repertoire of patients with periodontitis are oligoclonal compared with their peripheral blood, with a biased expression of TCR genes (23, 43–45).

Prior work comparing CD45RO expression in CD4⁺ and CD8⁺ T cells from healthy gingiva revealed that most CD4⁺ T cells are CD45RO⁺, indicating prior antigen experience (13). Our data aligns with those studies and challenges the notion of a purely quantitative increase in memory T cells during disease, since healthy gingiva is also densely populated with memory CD4⁺ T cells (46). Instead, the human data suggests qualitative reprogramming within an existent CD45RO⁺CD4⁺ pool toward a Tfh/ph-like, B-helper phenotype. Critically, the periodontitis-associated cluster in human was transcriptomically related to the expanded and convergent clonotype found in mouse recovery and recurrence, as both display a Tfh/Tph-like phenotype. Re-analysis of the human dataset showed that the Tph-like cluster (C9) was the sole disease-associated T cell variation at two resolution levels. Pseudobulk differential expression across all gingival T cells showed significantly higher expression of CXCR5, CXCL13, BCL6, and TOX2 in periodontitis than in health, a reliable signature of the Tph/Tfh B-helper program that is enriched in the C9 cluster (18). Consistent with this, differential abundance analysis identified C9 as the only disease-associated cluster, with all remaining clusters far from significance. Although the signal was modest on its own and did not reach significance using the most conservative blocked model, it confirms the same population that carries the Tph-like transcriptional program as periodontitis-associated in a methodologically complementary way.

Tph and Tfh CD4⁺ T cells are distinct populations that share core features of B-helper function. Both express CXCL13, PD-1, CD40L, ICOS, and MAF, and both act on B cells largely through IL-21 (18). Their principal difference is in localization. Tfh cells support naïve and memory B cells within germinal centers of secondary lymphoid organs, while Tph cells occupy peripheral tissues, blood, and tertiary lymphoid structures (TLSs) (17), act mainly on memory B cells, and characteristically lack the robust CXCR5 and BCL6 expression that defines Tfh cells. In inflamed tissue, Tph cells may further acquire Th1- and Tfh-like features in response to local cytokines, including T-bet expression, which we also detected in our data set (18, 47). The boundary between the Tfh and Tph is difficult to draw in tissue (18). Our data confirm this ambiguity: both the human and mouse clusters show mixed expression of Tfh- and Tph-associated genes, with an overall phenotype closer to Tph, as expected in gingiva. At the same time, CXCR5 and BCL6 were differentially expressed in human periodontitis. Consistent with this, recent spatial transcriptomic studies of human gingiva have identified TLSs in periodontitis, with a T cell compartment that includes follicular helper (Tfh) and follicular regulatory (Tfreg) cells expressing PD-1, CD69, and CD200 (48). We therefore used “Tfh/Tph-like” to describe a phenotypically mixed population rather than to assert a single lineage assignment.

Our findings raise new questions about the role of Tfh/Tph cells in recurrent periodontitis, particularly their influence on B cell function within the gingival tissue. As disease progresses and becomes chronic, B cells and plasma cells accumulate in the gingival mucosa and are central to periodontal immunopathology (49). Through antibody production, antigen presentation, cytokine secretion, and complement activation, they can either restrain or drive disease progression (50). Because B cell behavior depends heavily on the local microenvironment, Tfh/Tph-derived signals via the IL21/CXCL13 axis may be a key determinant of their pathogenic potential (51).

This study has limitations. Because each scRNA-seq and TCR-seq sample required tissue pooled from a large number of animals, the number of independent biological replicates was limited, restricting statistical power and the ability to assess biological variability. Consequently, findings from these datasets are primarily descriptive and exploratory and should be interpreted with caution. The model induces acute experimental depletion of the regulatory compartment, a more severe perturbation than physiological Treg fluctuations that occur over decades in patients. Also, the human scRNAseq data is a reanalysis of a published dataset and implies its cohort constraints (25). Nonetheless, to our knowledge, this is the first report of T cell clonal convergence in a model of recurrent mucosal inflammation.

Taken together, this study supports a paradigm in which clinical recovery does not ensure local T cell repertoire reset; instead, antigen-experienced, tissue-adapted CD4⁺ T cells accumulate and develop overtime a reservoir participating in disease progression. Immune regulatory disruptions may concurrently enhance the cumulative imprint on mucosal tissues. Clinically this model implies that each recovery phase post-therapy may involve the reshape of the patient’s susceptibility to recurrence, at least in part, by a combined history of previous disease events and transient regulatory perturbations during their lifespan.

## Materials and methods

### Mice

B6.129(Cg)-Foxp3^tm3(HBEGF/GFP)Ayr/J (Strain #016958) and B6.129S4-Il17a^tm1.1Lky/J (Strain #032406, SMART-17A) mice were obtained from The Jackson Laboratory and crossed to generate a line carrying both alleles (referred to as Foxp3^DTR/GFP^ in the article). The Il17a^Smart17A^ allele encodes a bicistronic hNGFR surface reporter for Il17a expression; hNGFR detection did not perform reliably, and this allele was not phenotyped in this study. The C57BL/6 congenic strain B6.SJL-Ptprca Pepcb/BoyJ (Strain #002014, Jackson Laboratory) was used as the host strain for adoptive transfer experiments. Mice were maintained under specific pathogen-free (SPF) conditions in standard animal facilities at the University at Buffalo and the ADA Forsyth Institute. Age-matched mice entered the model at 8–12 weeks of age; given the ∼50-day model duration, mice reached ∼18 weeks of age by the study endpoint. Genotyping was performed by Transnetyx® using their standard PCR-based service. Both male and female mice were included in all experiments, distributed across all experimental groups. All animal procedures were approved by the Institutional Animal Care and Use Committees (IACUC) of the University at Buffalo (protocol 202500003) and the ADA Forsyth Institute (protocols 19-001, 19-011, and 22-002).

### The Sensitized Recurrent Periodontitis Model (SeRPD)

The experimental model consisted of four phases: Baseline (BAS), Periodontitis (PD), Recovery (REC), and Recurrence (RCU). Baseline mice were age matched untouched mice. Periodontitis was induced in Foxp3 ^DTR/GFP^ mice by placing 5-0 silk ligatures around the maxillary second molars on day 0 for 10 days. Diphtheria toxin (DT; 50 µg/kg in sterile PBS) was administered intraperitoneally on days 1 and 2 to ensure transient depletion of the Treg cell population. The recovery phase began on day 10 with ligature removal, followed by a 30-day healing period before recurrence induction. Recurrence was subsequently induced by reapplying the ligatures for 10 days without further DT administration. This design included four experimental groups: Baseline (BAS; untreated control mice), Periodontitis (PD; euthanized on day 10 after the PD phase), Recovered (REC; euthanized on day 40), and Recurrence (RCU; euthanized on day 50). Unless otherwise specified, REC mice received DT as described above; in a subset of experiments, an RND group (ligature and recovery timeline identical, without DT administration) was included as indicated in the relevant figure legends and text.

### Gingiva Cell isolation for single-cell sequencing

Each group had two biological replicates, with each replicate containing at least 10,000 live CD45⁺ cells sorted from the gingiva of pooled mice per replicate: 9 mice for PD, 10 for RCU, and 18 for BAS and REC. Pooling was required because of low gingival immune cellularity and small tissue size (1 mm × 0.25 mm) which precludes assessment of inter-animal variability. Although the gingiva of recovered mice showed a high frequency of CD45⁺ cells within all live cells, these animals were no longer actively inflamed; consequently, the number of cells isolated per mouse was comparable to baseline, requiring additional animals per replicate to reach the 10,000-cell target. Before tissue isolation, we intracardially perfused all mice with PBS to remove peripheral blood. Gingival samples were placed into gentleMACS™ C Tubes (Miltenyi Biotec) containing 3 mL of enzymatic digestion medium composed of 2.7 mg/mL Collagenase IV (Gibco, cat# 17104019) and 0.15 µg/mL DNase I (STEMCELL, cat # 100-0762) in RPMI 1640. Tissue dissociation was performed using the gentleMACS™ Octo Dissociator with Heaters (Miltenyi Biotec) with a custom program to dissociate the tissue for 50 minutes at 37 °C. The resulting cell suspension was passed through a 70 µm cell strainer and washed with 35 mL of RPMI medium containing 1% penicillin–streptomycin and 0.15 µg/mL DNase I. Cells were then resuspended in complete media and stained with anti-mouse CD45 antibody (#103106, BioLegend) and 7AAD (eBioscience) for 15 min in the dark. At least 10,000 cells with >90% viability were sorted per sample using a FACS Aria II cell sorter (BD) at the Harvard Medical School Immunology Flow Cytometry Core.

### Gene Expression and TCR libraries preparation and sequencing

FACS-sorted CD45^+^ cells were subjected to the Chromium Single-Cell 5′ Reagent kits v2 workflow with feature barcode technology (10x Genomics) to generate a Gel Beads-in-emulsion (GEMs), where all generated cDNA shares a common 10x Barcode. Single-cell V(D)J libraries from amplified V(D)J amplified cDNA and single-cell 5’ gene expression libraries from amplified cDNA derived from poly-adenylated mRNA were generated at the Harvard Medical School Single Cell Core. The Gene Expression (GEX) and T cell receptor (TCR) libraries were then sequenced on a NovaSeq6000 system (100 cycles) (Illumina) at the University at Buffalo Genomics and Bioinformatics Core Facility.

### Alignment, quality control, and pairing of scRNA-seq and TCR seq data

Post-sequencing the data were processed by the Genomic and Bioinformatics Core at the University at Buffalo as follows: the raw data were converted to FASTQ format using BCL Convert (v4.4.4) with default parameters. The resulting FASTQ files were processed with the 10x Genomics Cell Ranger multi pipeline (v9.0.0), utilizing the 2024 GRCm39 reference genome and default settings, with the cell annotation model parameter set to auto to enable adaptive cell type annotation. The resulting gene expression (GEX) and T cell receptor (TCR) outputs were subsequently imported into R (v4.5.1 and analyzed using Seurat (v5.4.0) and scRepertoire (v2.6.2). The datasets were integrated into a single Seurat object via the combineTCR and combineExpression functions to ensure accurate mapping of clonotype information to individual transcriptomes. After integration, cells were filtered to retain those with at least 100 detected features and less than 20% mitochondrial gene content, thereby preserving transcripts with low abundance while excluding cells exhibiting elevated mitochondrial expression indicative of poor quality or apoptotic states.

### Gene expression and T cell antigen receptor analysis

To analyze T cell clonality in the model, cells were assigned to the same clonotype using a stringent criterion: (1) identical V and J gene usage and (2) an identical CDR3 nucleotide sequence for the paired chains (both α and β when available, or the single productive in-frame chain). Within each sample, clonotypes represented by ≥ 2 cells (distinct 10x barcodes) were classified as clonally expanded, whereas unique clonotypes were considered non-expanded. Because gingival immune cells were pooled from multiple mice per replicate, clonality was assessed at the pooled sample level, and inter-animal clonal variability could not be evaluated.

### Flow Cytometry analysis

Mice gingival tissues were collected and digested in RPMI containing collagenase type IV (2.7 mg/mL, Gibco) and DNase I (0.15 µg/mL, Stemcell) with 1% penicillin–streptomycin (Gibco) for 1 hour at 37 °C under gentle agitation every 10 min. Cervical lymph node single-cell suspensions were obtained by dissociating tissue over a 70 μm strainer using a syringe plunger, then washed with FACS buffer (PBS + 5% FBS). Gingival cell suspensions were filtered through a 70 μm strainer and washed with 35 mL of DNase Buffer (0.15 µg/mL DNase I + 1% penicillin–streptomycin in RPMI). Both gingiva and lymph node samples were stained with Zombie B550 Fixable Viability Kit (BioLegend) for 30 min at room temperature, then washed with FACS buffer. Because gingival tissue yields limited cells, we fixed gingival cells with 2% paraformaldehyde for 10 min at room temperature and washed with FACS buffer before Fc block and surface staining. CLN samples were processed directly from without fixation at this stage. Fc receptors were blocked with TrueStain FcX (anti-mouse CD16/32, BioLegend) for 10 min at room temperature, followed by a FACS buffer wash. Surface staining was performed with the antibodies listed in **Supplementary Table 5** for 30 min at 4 °C using Brilliant Stain Buffer (BD Horizon) per manufacturer instructions, followed by a final FACS buffer wash. Unstained and single-stained gingiva and lymph node samples, together with single-stained beads for low-abundance markers (AbC™ Total Antibody Compensation Bead Kit, Thermo Fisher), were used to perform spectral unmixing and extract autofluorescence in SpectroFlo (Cytek). Fluorescence-minus-one (FMO) controls were included for gating validation. Data was acquired on a Cytek Aurora at the Optical Imaging and Analysis Facility, University at Buffalo. Unmixed data were analyzed in FlowJo v10.10.1. The gating strategy is shown in **Supplementary Figure 1**. The RND cervical lymph nodes flow cytometry experiments were performed using a conventional Attune NxT Cytometer (The ADA Forsyth Institute). Compensation was performed using the beads indicated above. Due to the constrains of using a conventional cytometer, a simplified antibody panel was used including: CD44 (PercpCy5.5), CD62L (APC), TRCβ (APCCy7), CD4 (BV7110), Live/Dead Fixable Yellow, CD103 (BV421). Because the CLN panels differed between RND compared to REC, these experiments included their own controls (Baseline) for quantifications, using corresponding littermates. For all experiments, sample sizes ranged from n = 4–6 mice per group depending on the experiment (see figure legends for experiment-specific n). Differences in sample size within individual experiments reflected mouse availability or sample loss due to technical issues (i.e., ligature or sample loss).

### Adoptive transfer experiment

CD4^+^ effector memory T cells (TEM), gated as CD4^+^ CD44^+^ CD62L^−^ CD25^−^, were isolated from cervical lymph nodes (≈1×10⁶ cells) and spleen (≈4×10⁶ cells) of donor animals (Foxp3^DTR/GFP^, CD45.2) by fluorescence-activated cell sorting. Female and male mice were distributed across all groups for both host and donors; average donor age was 18 weeks, and average host age was 15.2 weeks at euthanasia. Single-cell suspensions were stained with CD4 (BV711, 1/200), CD25 (PE, 1/200), CD44 (BV785, 1/200), CD62L (Alexa Fluor 700, 1/200), and DAPI (1/1000) in FACS buffer (PBS + 5% FBS), and sorted using a BD FACSAria Fusion cell sorter (University at Buffalo Optical Imaging and Analysis Facility). Post-sort purity was confirmed by re-analysis, typically ≈90%. Sorted cells were washed with sterile PBS, and cell count and viability were assessed by Trypan blue exclusion using an automated cell counter (TC20, Biorad). Cells were resuspended at 5×10⁶ cells/100 µL sterile PBS and filtered through a 30 µm cell strainer prior to retro-orbital injection into isoflurane-anesthetized host mice (CD45.1). Two mice were pooled per replicate for the recovery groups (RND and REC), and 3–4 mice were pooled per replicate for the baseline (BAS) group, with n=6 biological replicates (pools) per group. 5-0 silk ligatures were applied to bilateral second maxillary molars of host mice, under isoflurane anesthesia, the same day of the adoptive transfer and left in place for 7 days until readout. Harvested maxillae were subsequently used for micro-CT and TRAP staining.

### TRAP staining

Osteoclasts were identified on paraffin-embedded, decalcified maxillary sections by tartrate-resistant acid phosphatase (TRAP) staining according to standard protocols. Briefly, FFPE sections (4–5 µm) were dried at 37°C overnight, then baked at 60°C for 1 hour before staining. Slides were deparaffinized in xylene (3 × 5 min) and rehydrated through a graded ethanol series (100% and 95% EtOH, 20 dips × 2 each) to dH₂O. TRAP enzyme reaction was performed by incubating sections in a working solution of 0.1 M acetate buffer (pH 5.0) containing 0.3 M sodium tartrate, Triton X-100, and Naphthol AS-MX phosphate (Sigma), pre-warmed to 37°C, with Fast Red Violet LB Salt (Sigma) added immediately before use. Slides were incubated at 37°C for 40 –60 min and monitored for red/magenta signal development on bone surfaces. Sections were counterstained with hematoxylin, dehydrated through ethanol and xylene, and mounted with Cytoseal 60 (Epredia). TRAP-positive osteoclasts were identified as multinucleated (≥3 nuclei) cells with bright red/magenta cytoplasmic staining located on or adjacent to bone surfaces. Images were acquired using a Nikon Ni-E upright microscope (Optical Imaging and Analysis Facility, University at Buffalo) and analyzed using ImageJ software.

### Human gingiva samples

Study participants were recruited from patients undergoing periodontal surgery, third molar tooth extraction, or orthodontic tooth extraction as part of their standard dental care at The ADA Forsyth Faculty Associates Clinic and the Center for Clinical and Translational Research. All participants were enrolled under the Institutional Review Board (IRB)–approved protocol IRB# 21-06 at The ADA Forsyth Institute and provided written informed consent prior to participation. Periodontal disease was defined according to the most recent 2017 World Workshop (52) as having at least two interproximal sites (on different teeth) with ≥5 mm probing depth and ≥3 mm clinical attachment loss. A healthy periodontal status was defined as probing depths ≤3 mm, no bleeding on probing, and absence of clinical signs of inflammation. Inclusion criteria required that each subject: (1) provided written informed consent, (2) was medically healthy based on medical history, and (3) had at least 16 natural teeth (excluding third molars). Exclusion criteria included: antibiotic use within one month prior to sample collection; systemic diseases or conditions interfering with tissue collection or analysis (e.g., autoimmune disease, diabetes); chronic use of medications such as steroids, NSAIDs (ibuprofen, aspirin) for more than 2–3 days per week, anticoagulants, or immunosuppressants; bleeding disorders; pregnancy or nursing; and cigarette smoking (including e-cigarettes or vaping) within one year prior to enrollment. Gingival samples containing gingival epithelium and underlying connective tissue were obtained by excisional incision based on the surgical treatment plan and standardized in size during trimming and embedding. All samples were fixed in 10% neutral-buffered formalin and embedded in paraffin (FFPE) for subsequent analyses.

### Imaging mass cytometry

Processing of human gingiva FFPE blocks, imaging and downstream analyses were performed at the Center for Cancer Biomarkers (CCBIO) at the University of Bergen. Imaging mass cytometry (IMC) was performed using the Hyperion imaging system (Standard Biotools) according to the manufacturer’s protocol. The IMC antibody panel was adapted from a previously developed and validated multiplex panel described by Tornaas et al. (53), with modifications to accommodate the specific objectives of the present study. The panel was designed to characterize immune cell populations in gingival tissue, with particular emphasis on T-cell subsets and antigen-experienced T cells. The antibody staining procedure was based on the previously published protocol (Tornaas et al., 2024), with adaptations for gingival tissue. Briefly, regions of interest (ROI) were selected from 7 samples, and the areas were laser-ablated at 200 Hz, producing a pixel size/resolution of 1 μm^2^. Acquisition was done using CyTOF software v7.0 (Standard Biotools). Visualization and evaluation of high-quality images with selected combinations of markers were done using MCD Viewer v1.0.560.6 (Standard Biotools). Segmentation of raw IMC data was done following the Steinbock framework with a Docker container (Deepcell). Spillover correction of IMC data was done following the protocol established by the Bodenmiller laboratory. Final image analysis and quantification were performed using the TIFF files with Qupath and Fiji software, with support of the Optical Imaging and Analysis Facility, University at Buffalo.

### Re-analysis of published scRNAseq human data

Publicly available single-cell RNA-seq data were obtained from the Gene Expression Omnibus (accession GSE164241) (25). Raw reads for gingival mucosa samples from healthy donors (GM; n = 13) and patients with periodontitis (PD; n = 8) were downloaded from the Sequence Read Archive using fasterq-dump (SRA Toolkit v3.0.5); the healthy buccal mucosa samples (n = 8) were excluded. Read quality was assessed with FastQC (v0.11.9), and samples were screened for contamination using FastQ Screen (v0.14.1). Results were summarized with MultiQC (v1.31). Sequencing runs were pooled by sample and processed using Cell Ranger (v10.0.0; GRCh38-2024-A), yielding 21 samples from 18 donors (including separate repeat biopsies for PD164 [n = 3] and PD161 [n = 2]). Cells were filtered to retain those with 200–6,000 detected genes and <20% mitochondrial reads. Following batch correction with Harmony, cells were clustered using Seurat (R v4.2.0, Seurat v5.1.0, harmony v0.1.0). Total immune cells, and subsequently T cells, were sequentially isolated, re-integrated, and re-clustered. We excluded T/plasma-cell doublets (>10% immunoglobulin reads) and a ribosomal-gene-dominated cluster lacking lineage markers, resulting in a final dataset of 20,802 T cells across nine clusters (C1–C9). Differential abundance was evaluated via propeller (speckle) using logit-transformed per-sample proportions with donor blocking (within-donor correlation ρ = 0.73). Differential expression was analyzed on per-sample pseudobulk profiles using edgeR::voomLmFit with sample quality weights and donor blocking. P-values were adjusted using the Benjamini–Hochberg false discovery rate (FDR) method. Statistical workflows were executed in R v4.5.1 (limma v3.66.0, edgeR v4.8.2, speckle v1.10.0).

### Bone loss measurements

*Micro-computed tomography* (µCT): Maxillae were fixed in 10% neutral-buffered formalin at 4°C overnight and transferred to 70% ethanol prior to scanning using a Scanco µCT 100 system (Scanco Medical) at a resolution of 8 µm. Resulting DICOM files were analyzed with AnalyzePro (version 14, AnalyzeDirect) to quantify total and bone volume within defined anatomical landmarks, with normalization performed by using a fixed anatomical region of interest across samples. For µCT analysis, a subset of samples was excluded due to bone breakage during processing or scanning artifacts; these losses were unrelated to experimental condition. Final sample sizes per group are indicated in the corresponding figure legends.

*Stereomicroscopic morphometric analysis*: In separate experiments, maxillae were defleshed by mechanical removal of soft tissue and bleached with 10% hydrogen peroxide to expose the bone surface. Images were acquired using a stereomicroscope at 4X magnification, and the area between the cementoenamel junction and alveolar bone crest was measured using ImageJ software. In BAS, REC and RND comparison, a single baseline group was used since all samples were processed equally, unlike the flow cytometry experiment.

### Statistical Analysis

Data visualization and statistical analyses of scRNA-seq and TCR-seq data were performed using SAS version 9.4 and R version 4.4.3 in RStudio version 2024.12.1+563. Gene repertoires overlap between REC and RCU was assessed using the Jaccard similarity coefficient, with subsampling analyses conducted to evaluate robustness. Gene prevalence was summarized as the proportion of cells in which each gene was observed, and associations in gene prevalence between conditions were assessed using correlation analysis. Cell-type–specific differences were evaluated using pairwise comparisons of gene prevalence, with genes ranked by absolute differences in prevalence. Statistical analyses and visualization for all other data were performed using GraphPad Prism 10. Specific statistical tests used for each comparison are indicated in the results section and/or corresponding figure legends.

## Data availability

The mouse gingiva scRNA-seq and TCR data presented in this article are available from NCBI Gene Expression Omnibus (GEO) under the accession number GSE350875. The remaining data are available from the corresponding author upon reasonable request.

## Acknowledgements

The authors thank Andrew McCall, Director of the Optical Imaging and Analysis Facility at the University at Buffalo, for assistance with imaging analysis, and laboratory technician Lesli Nelson for technical support. We also gratefully acknowledge Tyler Chinsky from the University at Buffalo Genomics and Bioinformatics Core, the Center for Computational Research, and the Biostatistics Center for Collaborative Research and Data Coordination. This work was supported by the National Institute of Dental and Craniofacial Research (K99DE032704 and R00DE032704 to C. Alvarez Rivas), the Philips Oral Healthcare Young Investigator Research Grant (to C. Alvarez Rivas), the ADA Forsyth Pilot Grant (to B. Bor and C. Alvarez Rivas), and by the Trond Mohn Foundation (TMS2021STG03 to S. Suliman).

## Author contributions

S.A.: Formal analysis, Software

O.V.: Investigation

R.L.: Formal analysis, Methodology

S.S.: Methodology, Funding acquisition

M.S. and L.K.: Investigation

C.D.K.: Investigation, Formal analysis

A.H.A.: Formal analysis, Investigation

A.K., D.J., and B.C.: Investigation

Y.N.: Methodology

T.S., C.D., and N.D.H.: Investigation, Formal analysis

G.Z. and O.F.: Investigation

B.B.: Funding acquisition

S.Ah.: Formal analysis

H.H.: Resources

H.G., K.K., and K.A.: Conceptualization, review & editing

C.A.R.: Conceptualization, Methodology, Funding acquisition, Investigation, Writing

## Declaration of generative AI and AI-assisted technologies in the manuscript preparation process

During the preparation of this work, the authors used Claude Sonnet 5 (Anthropic) to assist with early data analysis tasks, including code development and initial exploration of cell-type annotations, as well as to detect grammar and typographical errors in the manuscript text. All final annotations were manually curated, against published literature and reference atlases, and all analytical decisions and interpretations were made independently by the authors. After using this service, the authors reviewed and edited the content as needed and take full responsibility for the content of the published article.

---

**Supplementary Figure 1.**
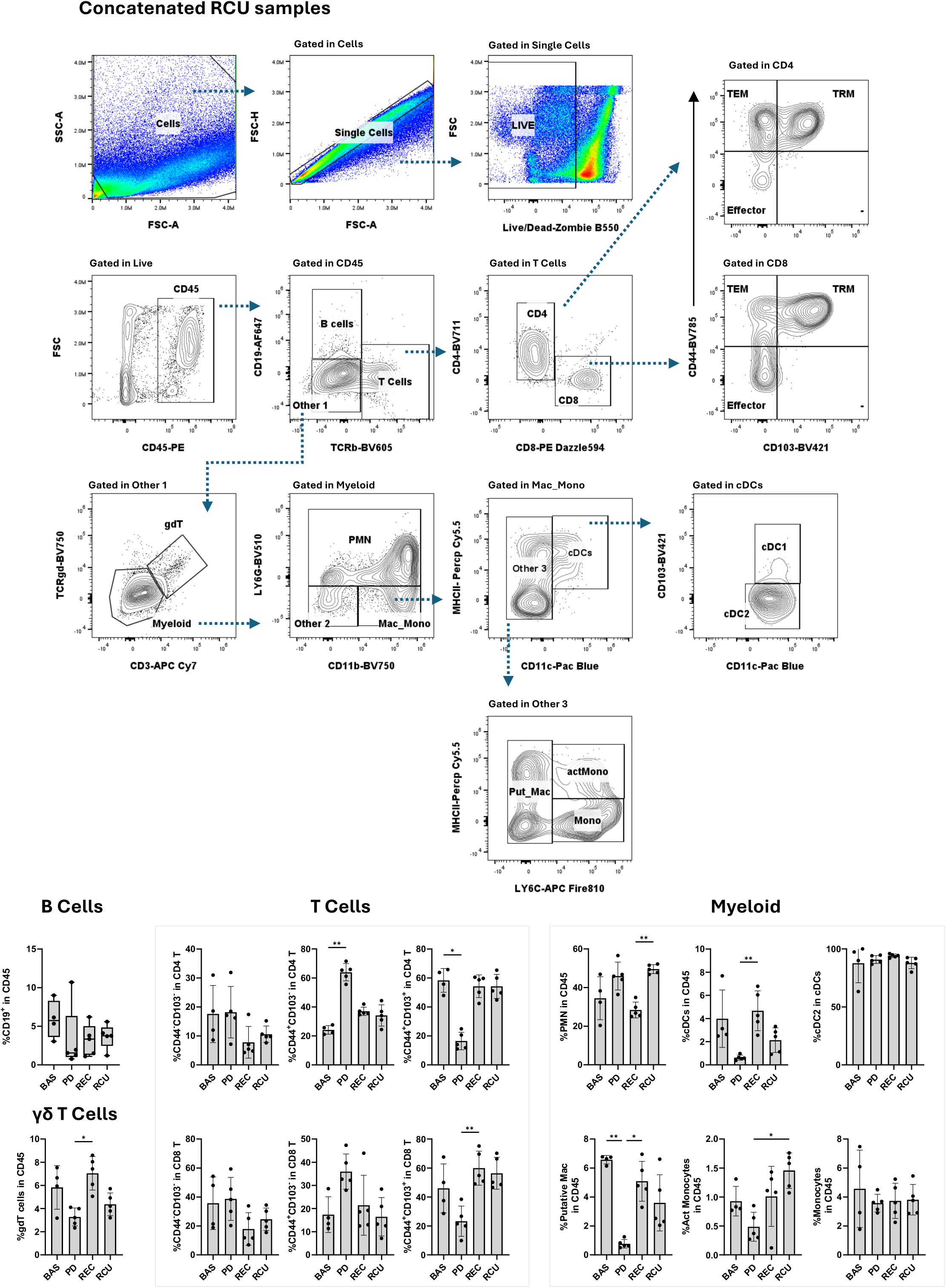
The SeRPD mouse model drives remodeling of the gingival immune landscape across disease phases. A) Representative gating strategy identifying CD45⁺ leukocytes, TCRβ⁺ T cells, CD4⁺ and CD8⁺ T cells, and myeloid populations in gingival tissue, shown for four concatenated Recurrent (RCU) samples. B) Quantification of frequencies of CD45⁺, TCRβ⁺CD4⁺, TCRβ⁺CD8⁺, and myeloid cell populations in gingival tissue per condition (BAS: n=4; PD, REC, and RCU: n=5 per group). Statistical comparisons performed using the Kruskal–Wallis test with post hoc correction for multiple comparisons; ns = p > 0.05, * p < 0.05, ** p < 0.01.

**Supplementary Figure 2.**
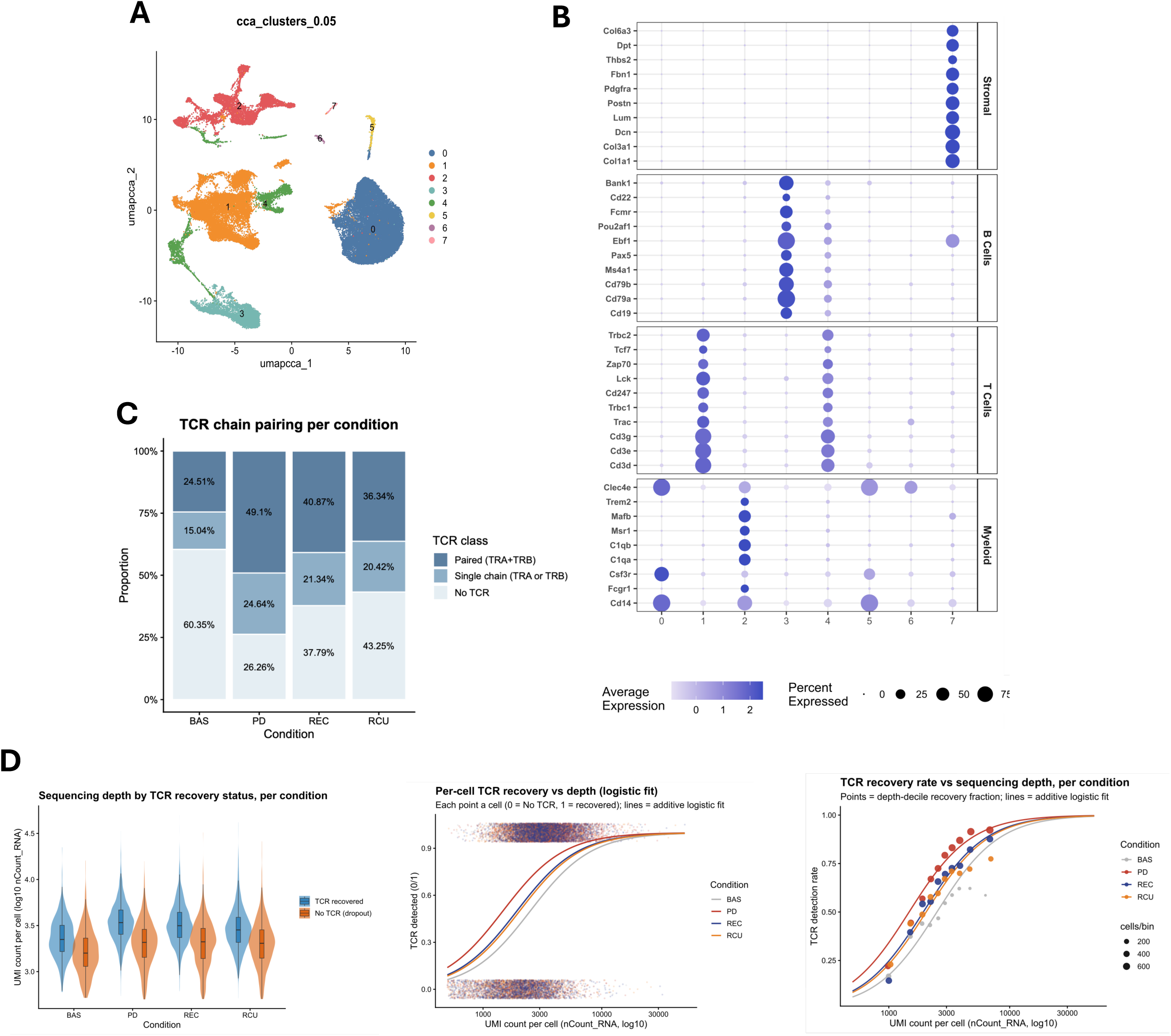
Global clustering of gingival immune cells, TCR chain recovery, and dependence of TCR recovery on sequencing depth. A A) UMAP of 51,572 gingival cells from all conditions after CCA integration, showing eight clusters (0–7) identified by unsupervised clustering. B) Dot plot of canonical marker genes across the eight clusters, grouped by major cell type: myeloid (clusters 0, 2, 5 and 6), T cells (1 and 4), B cells (3) and stromal cells (7). These annotations define the four populations shown in Figure 4B. Dot color represents scaled average expression; dot size represents the percentage of cells expressing each gene. C) Proportion of T cells with a paired (TRA + TRB), single-chain (TRA or TRB) or no detected TCR in each condition, among T cells retained after removal of non-T contaminants (n = 14,530). D) TCR recovery as a function of sequencing depth (UMI count per cell). Left: UMI count of cells with and without a recovered TCR in each condition. Middle: recovery status of individual cells (1 = recovered, 0 = no TCR) against UMI count, with logistic regression fits per condition. Right: TCR recovery rate within UMI-count deciles per condition; point size indicates the number of cells per decile, and lines show logistic regression fits. TCR recovery increased with sequencing depth independent of condition (logistic regression adjusted for condition: odds ratio 49 per 10-fold increase in UMI count, 95% CI 41–59; mixed model with sample as random effect: odds ratio 46, 95% CI 38–56; p < 2 × 10⁻¹⁶).

**Supplementary Figure 3.**
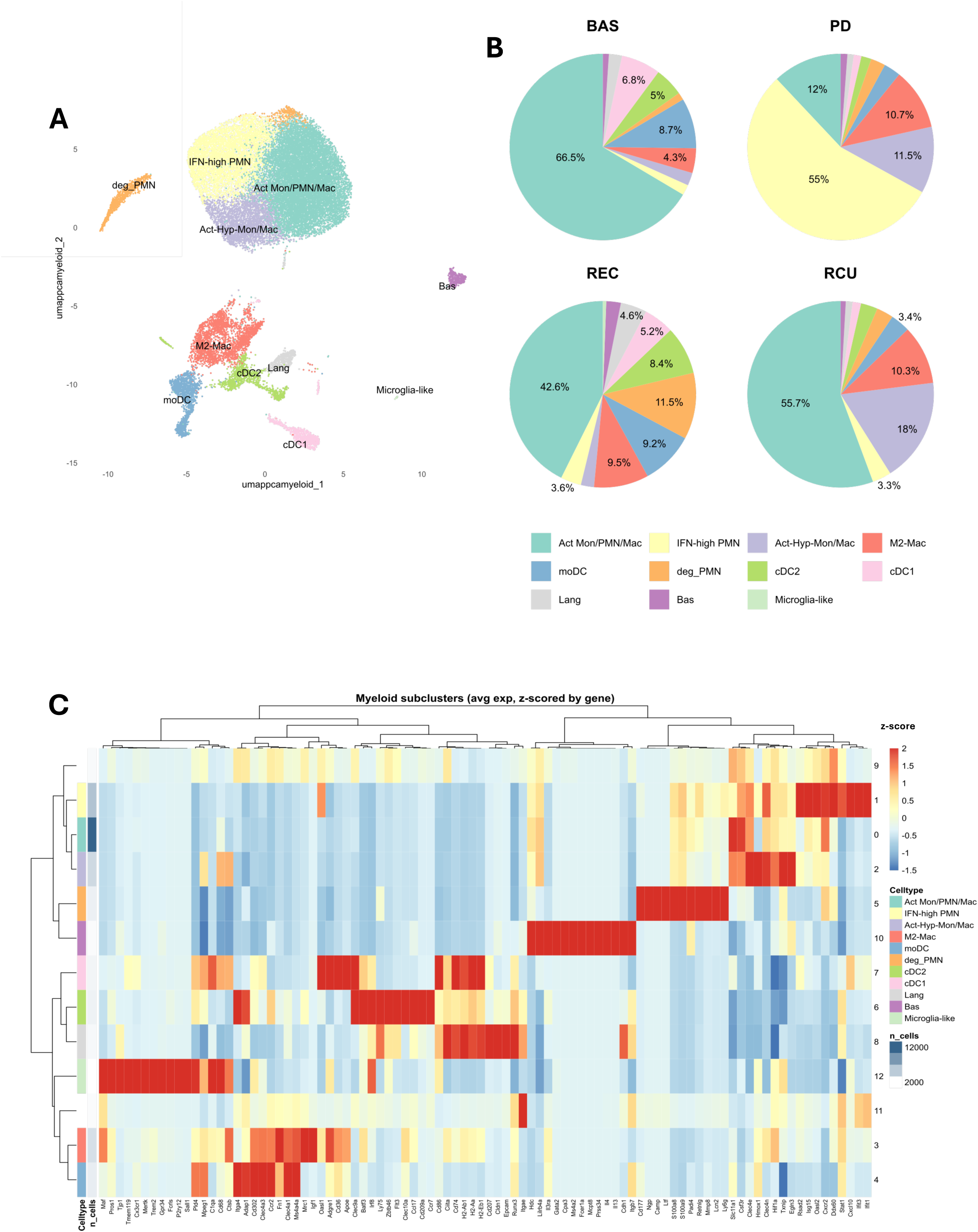
Myeloid subpopulations in gingiva across disease phases. A) UMAP of 27,740 gingival myeloid cells after sub-clustering, annotated into 11 subsets: activated monocytes/neutrophils/macrophages (Act Mon/PMN/Mac), interferon-high neutrophils (IFN-high PMN), activated hypoxia-adapted monocytes/macrophages (Act-Hyp-Mon/Mac), M2-like macrophages (M2-Mac), monocyte-derived dendritic cells (moDC), degranulating neutrophils (deg_PMN), conventional type 2 and type 1 dendritic cells (cDC2, cDC1), Langerhans cells (Lang), basophils (Bas) and microglia-like cells. B) Pie charts showing the proportion of each myeloid subset within each condition (BAS: 5,006 cells; PD: 7,328; REC: 4,917; RCU: 10,489). C) Heatmap of canonical myeloid marker genes across all 13 myeloid subclusters (0–12) identified before filtering. Fill shows average expression per subcluster, z-scored per gene. Rows and columns are clustered hierarchically. Row annotations show the assigned subset and the number of cells per subcluster. Subclusters 9 (B-cell and high-mitochondrial signature) and 11 (T-cell signature, likely doublets) were excluded from (A) and (B).

**Supplementary Figure 4.**
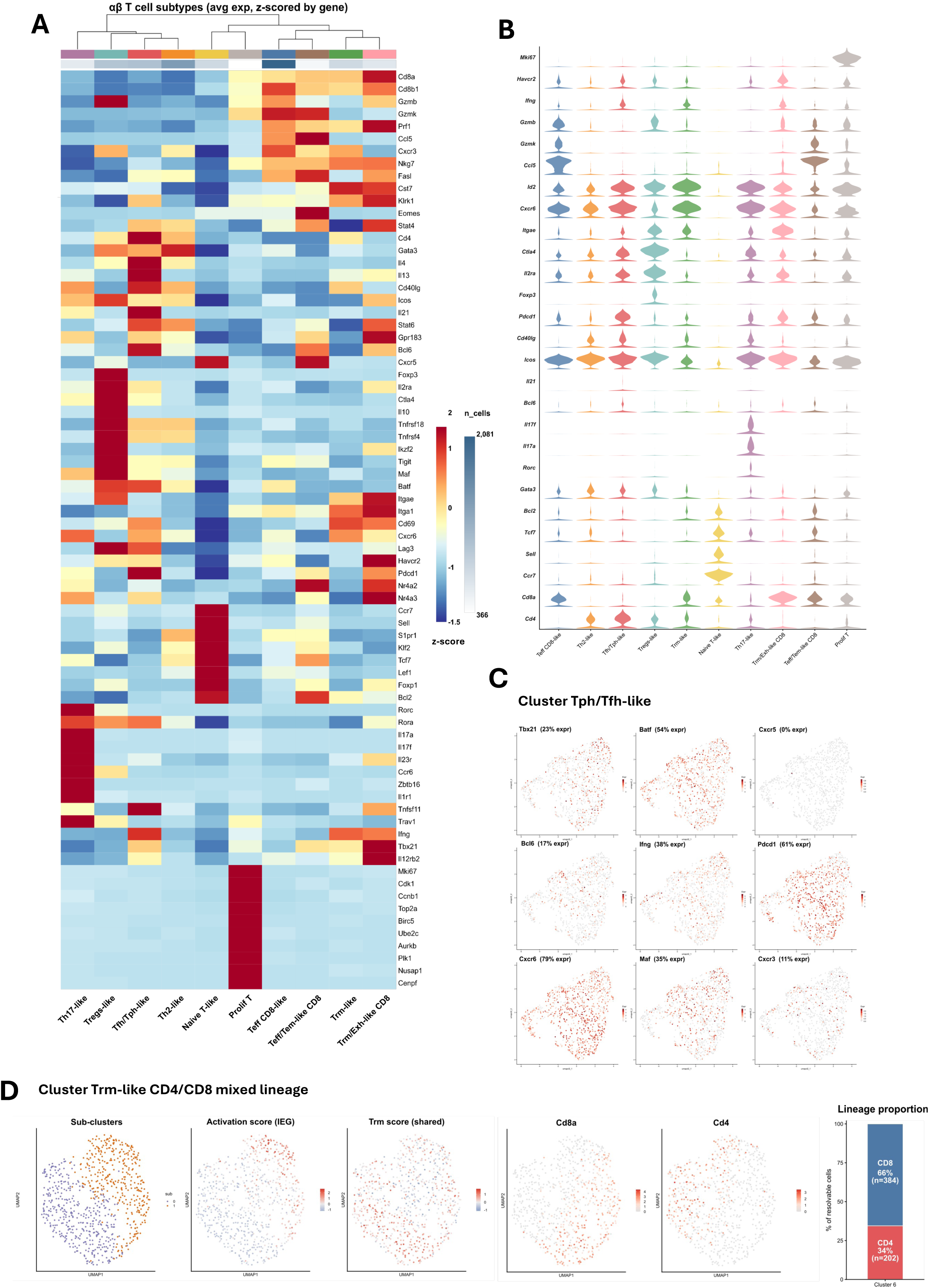
Annotation of αβ TCR⁺ T cell (single and paired chains) subclusters. A) Heatmap of lineage, transcription factor and functional marker genes across the ten αβ TCR⁺ T cell subclusters (n = 9,224 cells). Fill shows average expression per subcluster, z-scored per gene. Subclusters (columns) are clustered hierarchically, and the top annotation bar shows the number of cells per subcluster. B) Stacked violin plots of normalized expression of canonical and functional marker genes across subclusters. C) Expression of Tfh/Tph- and Th1-associated genes within the Tfh/Tph-like subcluster (n = 1,045 cells), shown on a UMAP re-computed for this subcluster alone. The percentage of cells expressing each gene is given in parentheses. D) Characterization of the Trm-like CD4/CD8 mixed-lineage subcluster (n = 781 cells) on a UMAP re-computed for this subcluster alone. From left: the two sub-clusters identified by unsupervised re-clustering; module scores for immediate-early activation genes (Atf3, Fosb, Dusp1, Nr4a1, Fos, Zfp36) and tissue-resident memory genes (Itgae, Itga1, Cxcr6, Id2); expression of Cd8a and Cd4; and the proportion of CD8⁺ (Cd8a⁺ or Cd8b1⁺, Cd4⁻; 66%, n = 384) and CD4⁺ (Cd4⁺, Cd8a⁻Cd8b1⁻; 34%, n = 202) cells among the 586 cells with a detectable lineage marker.

**Supplementary Table 1:** Clinical Metadata of Imaging Mass Cytometry samples.

| **Sample Code** | **Group** | **Site** | **Age** | **Sex** | **Race Ethnicity** |
|---|---|---|---|---|---|
| 11b | Periodontal Disease | Mandibular Molar | 28 | Female | White, Non-Hispanic or Latino |
| 6 | Periodontal Disease | Mandibular Molar | 62 | Male | White, Non-Hispanic or Latino |
| 1 | Periodontal Disease | Mandibular Molar | 21 | Male | White, Non-Hispanic or Latino |
| 5 | Periodontal Disease | Mandibular Molar | 35 | Male | White, Non-Hispanic or Latino |
| 7B | Healthy Tissue | Mandibular Molar | 27 | Female | White, Non-Hispanic or Latino |
| 10A/B | Healthy Tissue | Mandibular Molar | 62 | Male | White, Non-Hispanic or Latino |
| 8 | Healthy Tissue | Maxillary Molar | 72 | Female | White, Non-Hispanic or Latino |

**Supplementary Table 2:**
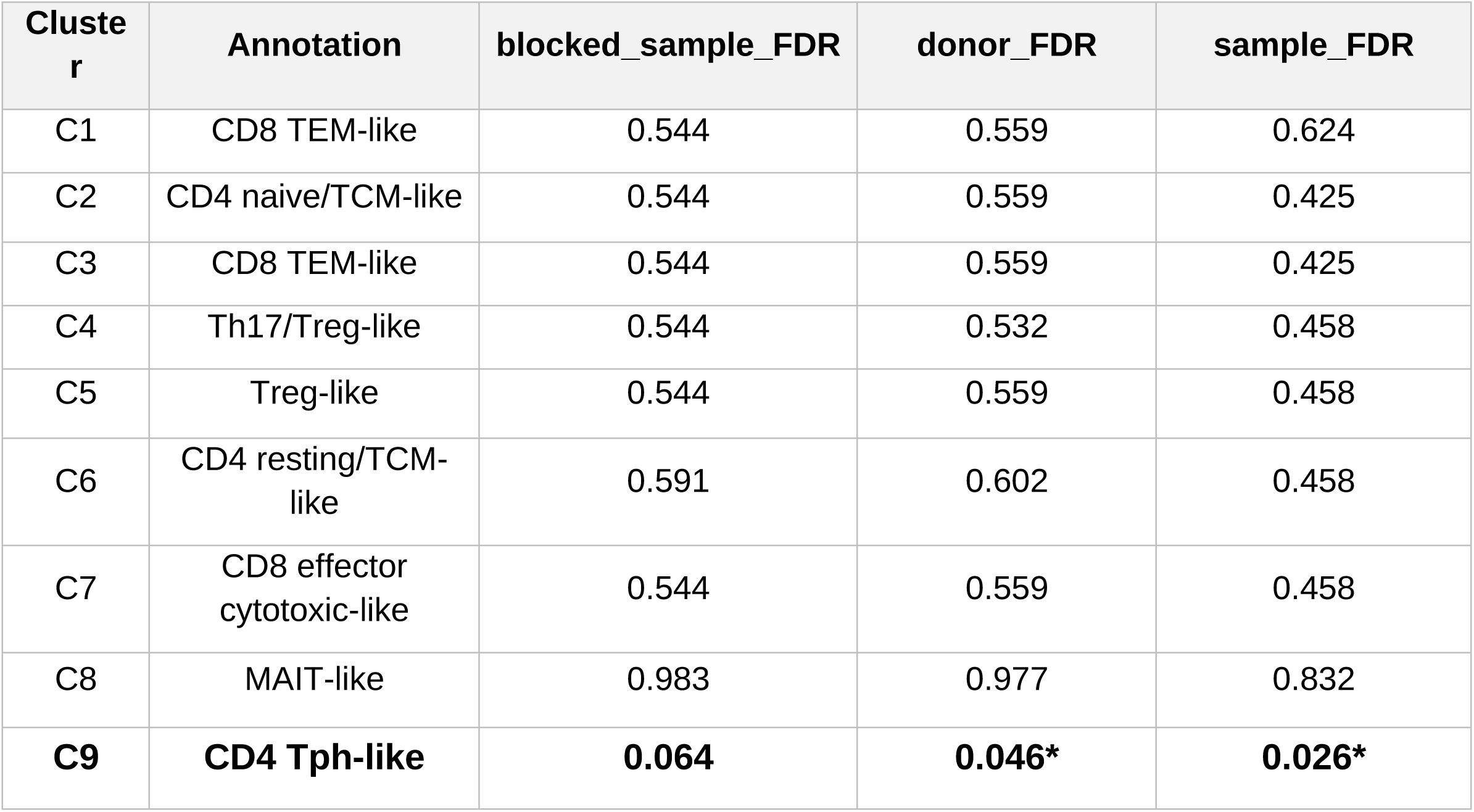
Human T cluster abundance was assessed using propeller, which models compositional cell-type proportions; we report results at the sample, donor, and donor-blocked levels as an internal robustness check across levels of aggregation.

**Supplementary Table 3:**
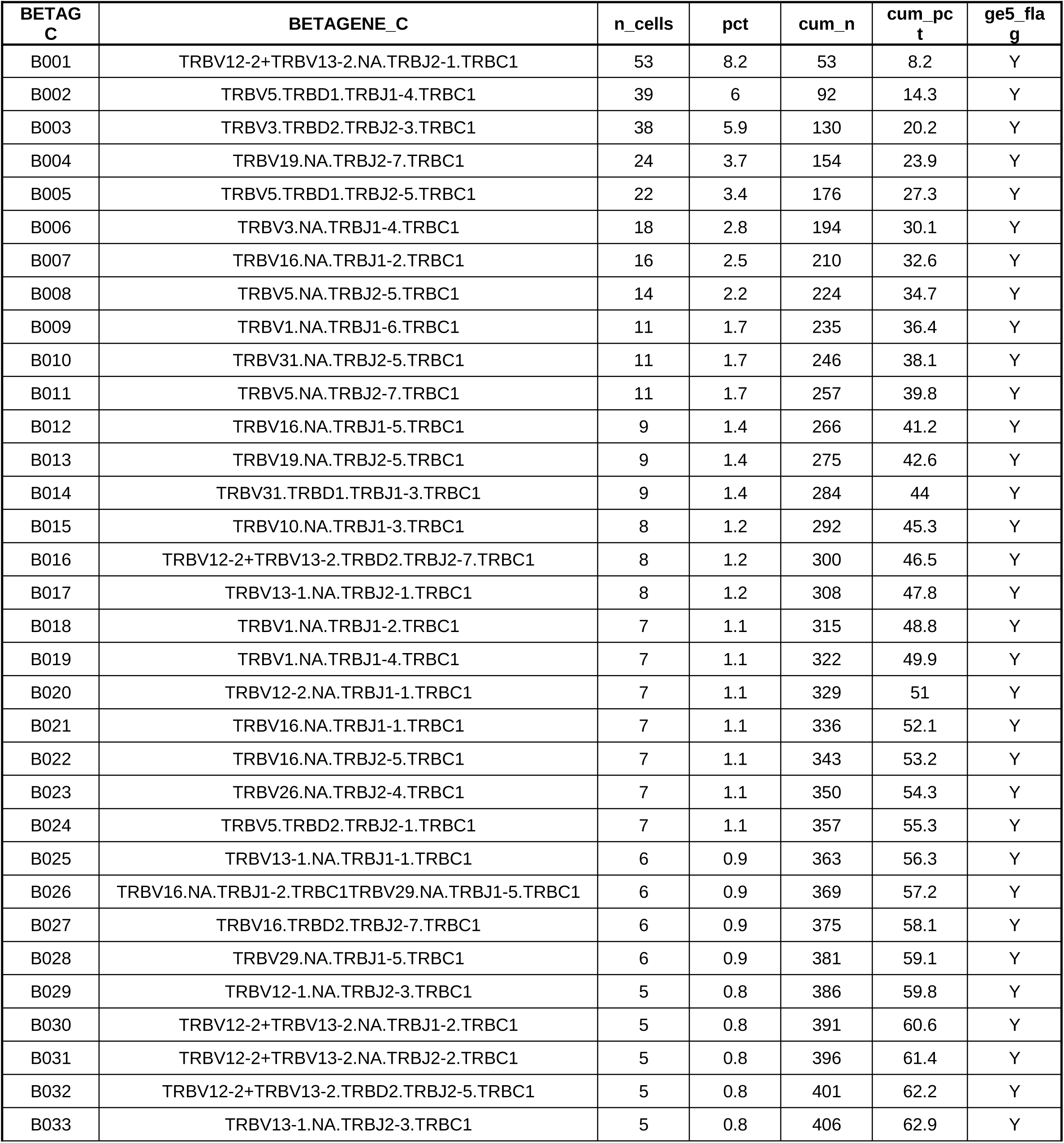
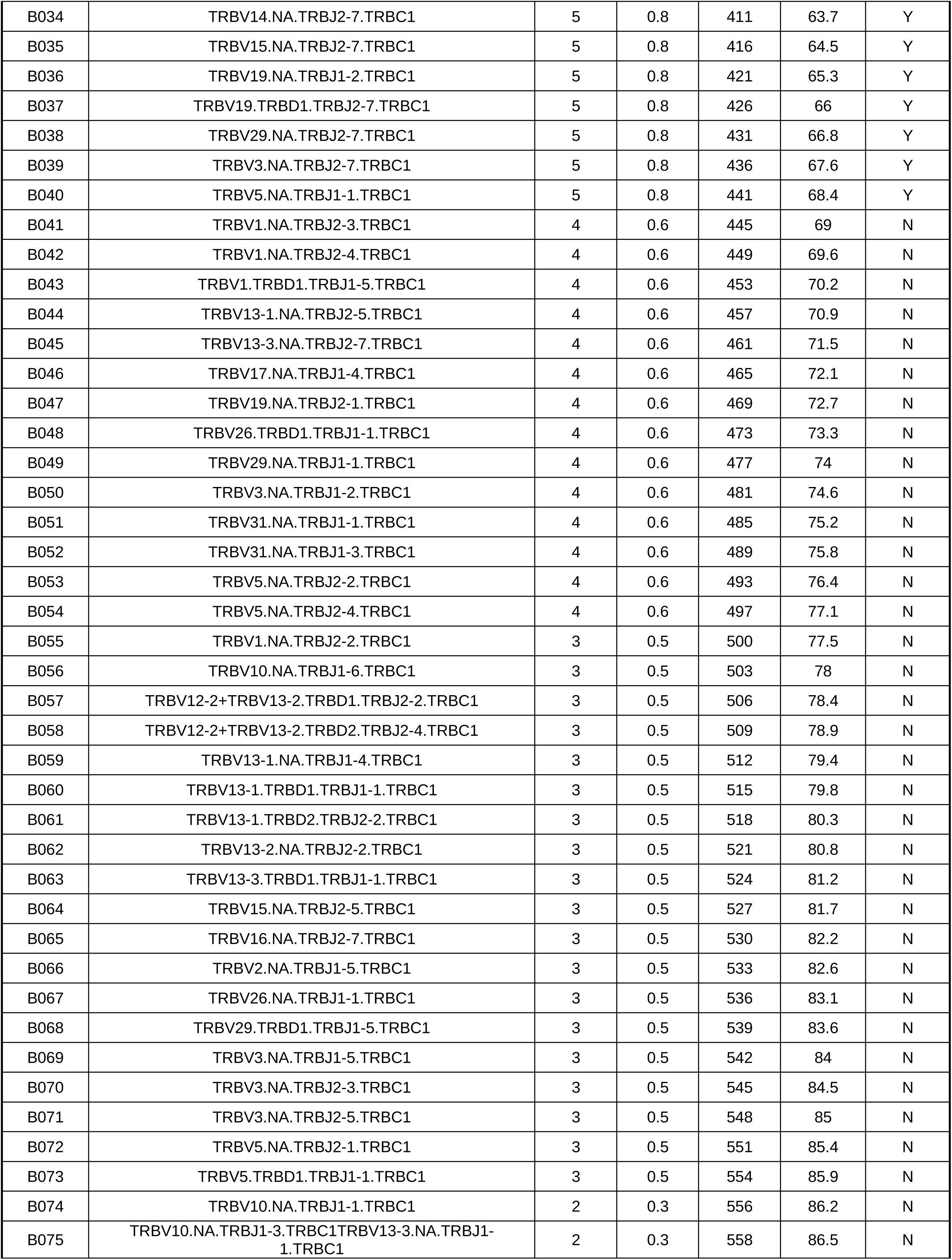
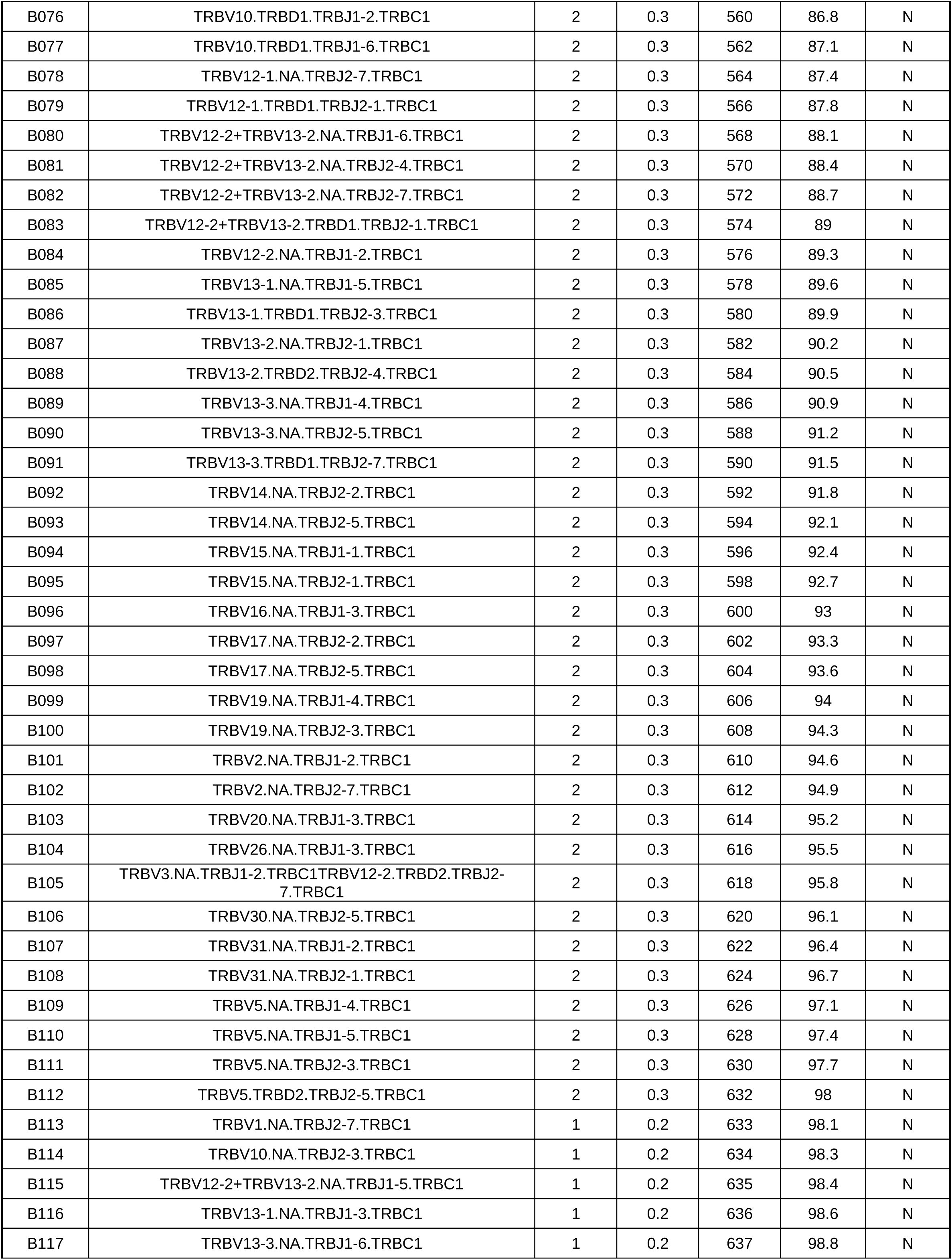
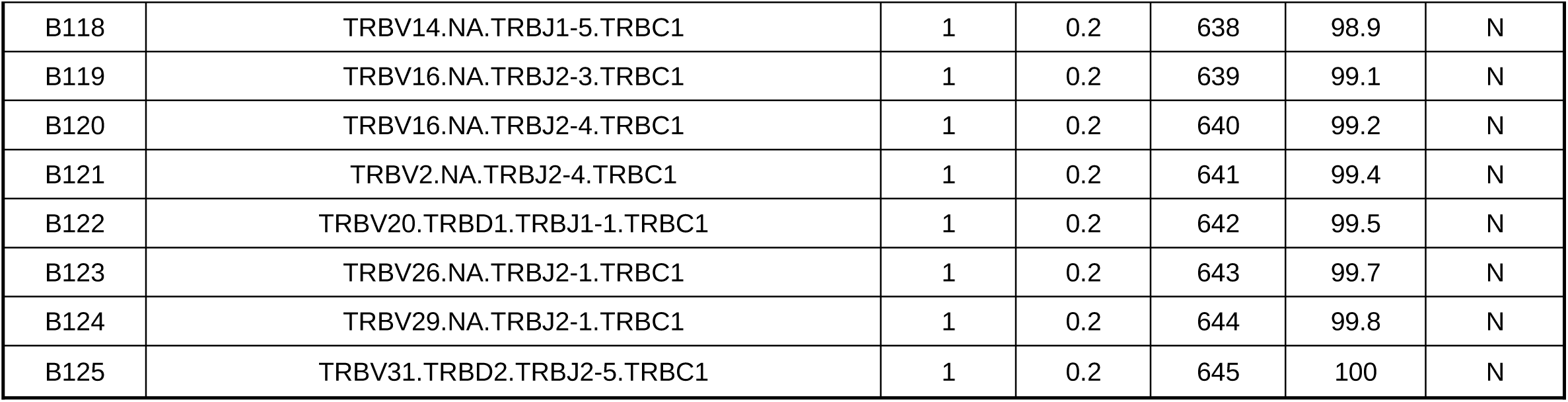
125 beta chain IDs (BETAGC = B001, B002,…) detected in REC and RCU expanded CD4^+^ T cells. Each ID comes with its beta gene string (BETAGENE_C, e.g. B001 = TRBV12-2+TRBV13-2.NA.TRBJ2-1.TRBC1), cell count (n cells), percentage (pct), cumulative number (cum n), cumulative percentage (cum pct), and a flag for beta chains with 5 or more identical cells (ge5_flag; Y:yes, N:no).

**Supplementary Table 4:**
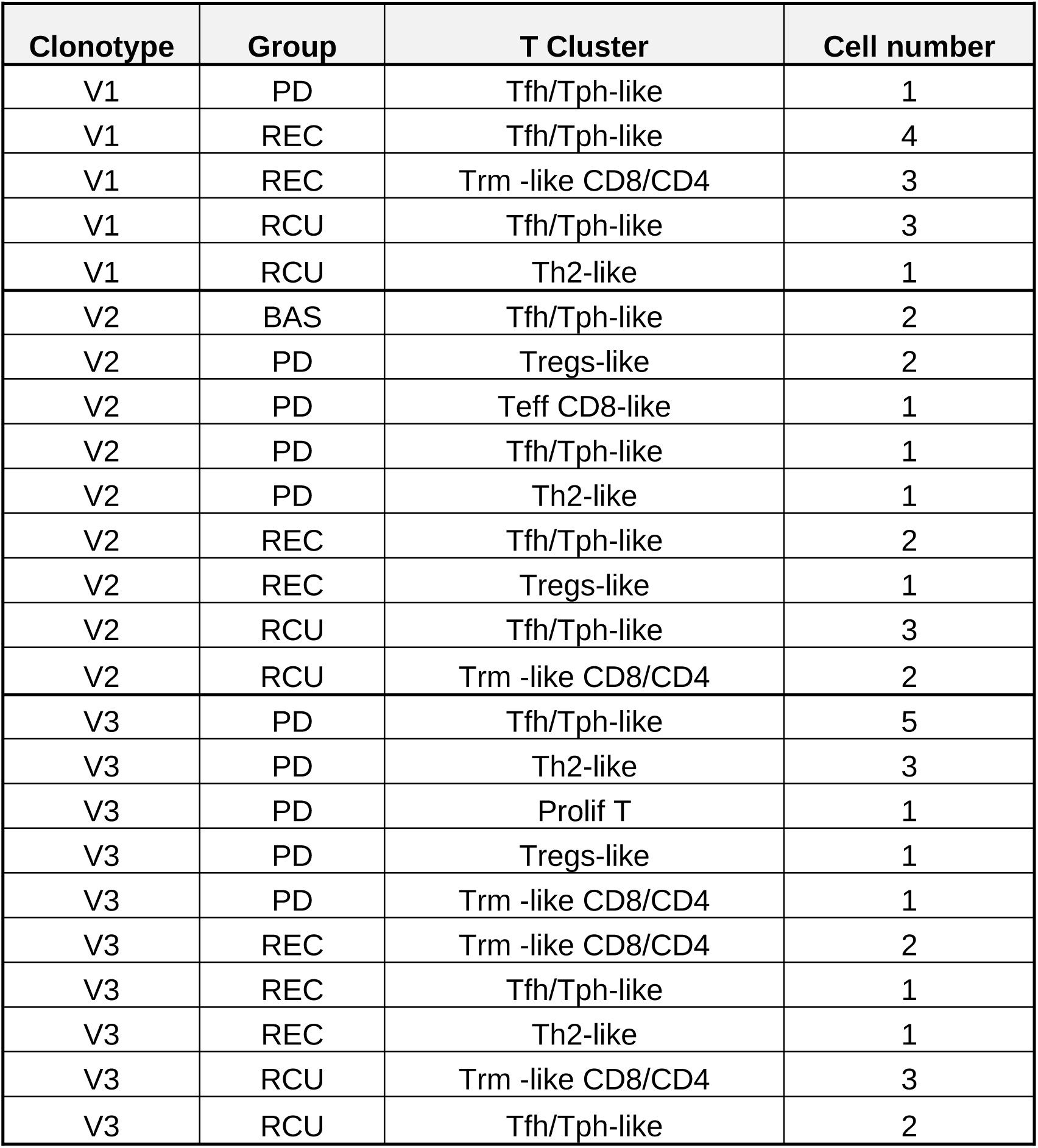
Number of cells per clonotype variant (V1-V3) and their mapping in each experimental groups and T cell clusters.

| Clonotype | Group | T Cluster | Cell number |
|---|---|---|---|
| V1 | PD | Tfh/Tph-like | 1 |
| V1 | REC | Tfh/Tph-like | 4 |
| V1 | REC | Trm -like CD8/CD4 | 3 |
| V1 | RCU | Tfh/Tph-like | 3 |
| V1 | RCU | Th2-like | 1 |
| V2 | BAS | Tfh/Tph-like | 2 |
| V2 | PD | Tregs-like | 2 |
| V2 | PD | Teff CD8-like | 1 |
| V2 | PD | Tfh/Tph-like | 1 |
| V2 | PD | Th2-like | 1 |
| V2 | REC | Tfh/Tph-like | 2 |
| V2 | REC | Tregs-like | 1 |
| V2 | RCU | Tfh/Tph-like | 3 |
| V2 | RCU | Trm -like CD8/CD4 | 2 |
| V3 | PD | Tfh/Tph-like | 5 |
| V3 | PD | Th2-like | 3 |
| V3 | PD | Prolif T | 1 |
| V3 | PD | Tregs-like | 1 |
| V3 | PD | Trm -like CD8/CD4 | 1 |
| V3 | REC | Trm -like CD8/CD4 | 2 |
| V3 | REC | Tfh/Tph-like | 1 |
| V3 | REC | Th2-like | 1 |
| V3 | RCU | Trm -like CD8/CD4 | 3 |
| V3 | RCU | Tfh/Tph-like | 2 |

**Supplementary Table 5:**
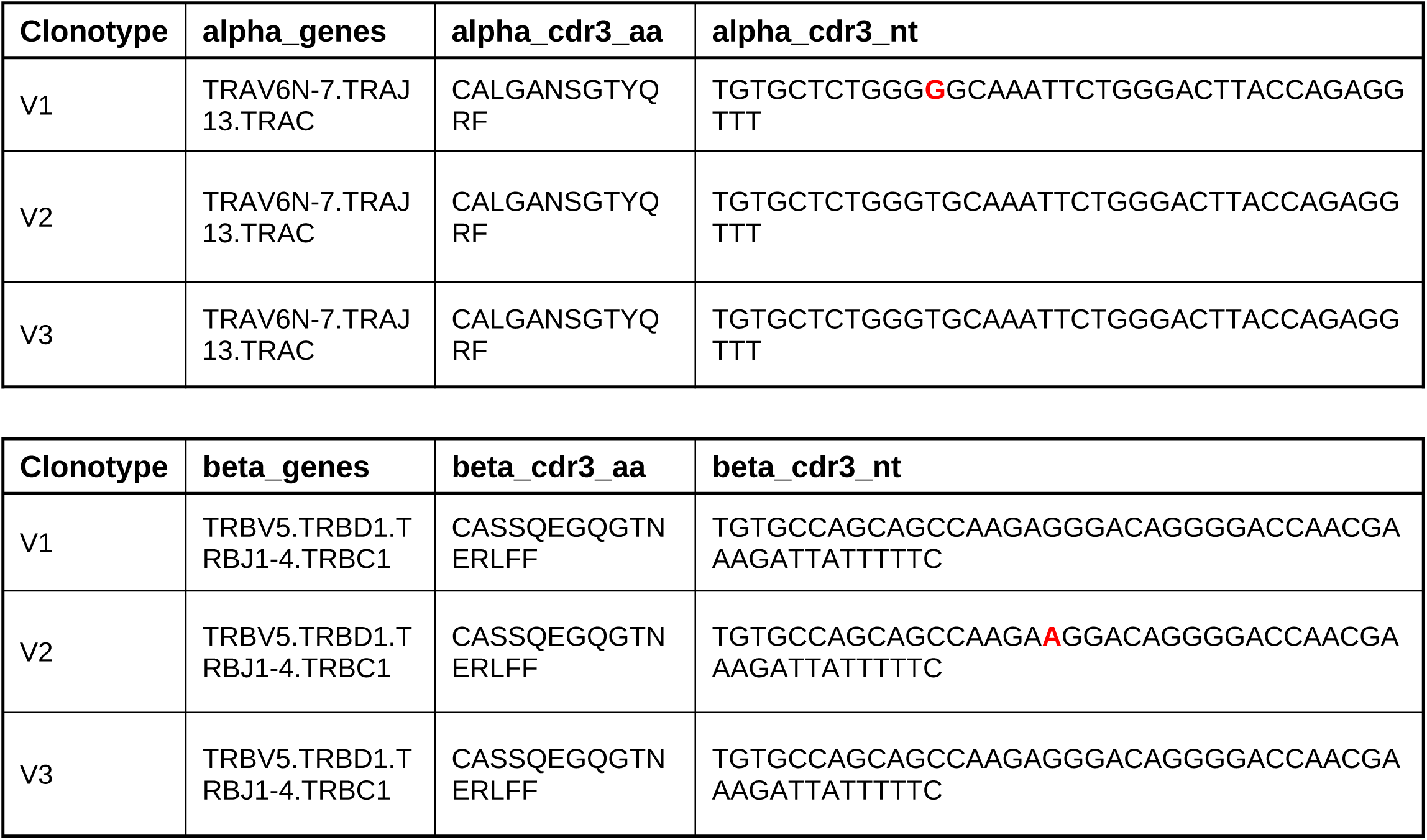
Convergent clonotypes variants of the same αβTCR protein. In red, identified unique nucleotides per variant.

**Supplementary Table 5:** Antibodies and dyes used in Flow cytometry and cell sorting experiments.

| Antibody | Clone | Cat Number | Supplier |
|---|---|---|---|
| PE anti-mouse CD45 | 30-F11 | 103106 | Biolegend |
| Brillant Violet 605 anti-mouse TCR β chain | H57-597 | 109241 | Biolegend |
| APC/Cyanine7 anti-mouse TCR β chain | H57-597 | 109219 | Biolegend |
| Brilliant Violet 711 anti-mouse CD4 | GK1.5 | 100447 | Biolegend |
| PE Dazzle 594 anti-mouse CD8a | 53-6.7 | 100761 | Biolegend |
| Brillant Violet 421 anti-mouse CD103 | 2E7 | 121421 | Biolegend |
| Brillant Violet 785 anti-mouse/human CD44 | IM7 | 103041 | Biolegend |
| PerCP/Cyanine5.5 anti-mouse/human CD44 | IM7 | 103031 | Biolegend |
| Alexa Fluor 700 anti-mouse CD62L | MEL-14 | 104426 | Biolegend |
| APC anti-mouse CD62L | MEL-14 | 104411 | Biolegend |
| Brillant Violet 650 anti-mouse TCRγδ | GL3 | 118147 | Biolegend |
| APC/ Cyanine 7 anti-mouse CD3 | 145-2C11 | 100329 | Biolegend |
| Alexa Fluor 647 anti-mouse CD19 | 6D5 | 115525 | Biolegend |
| Brillant Violet 750 anti-mouse/human CD11b | M1/70 | 101267 | Biolegend |
| Pacific blue anti-mouse CD11c | N418 | 117321 | Biolegend |
| APC/Fire 810 anti-mouse LY-6C | HK1.4 | 128055 | Biolegend |
| Brillant Violet 510 anti-mouse LY-6G | 1A8 | 127633 | Biolegend |
| PerCP/Cyanine 5.5 anti-mouse I-Ab | AF6-120.1 | 116415 | Biolegend |
| PE/Fire 810 anti-mouse CD138 (Syndecan-1) | 281-2 | 142545 | Biolegend |
| Zombie B550 Fixable Viability Kit | - | 423108 | Biolegend |
| LIVE/DEAD™ Fixable Yellow Dead Cell Stain Kit | - | L34959 | Thermo Fisher |
| DAPI | - | D3571 | Thermo Fisher |
| 7 AAD | - | 50-112-8859 | eBioscience |

